# Glc7/PP1 triggers Paf1 complex dissociation from RNA polymerase II to enable transcription termination

**DOI:** 10.1101/2025.09.03.674035

**Authors:** Sanchirmaa Namjilsuren, Karen M. Arndt

## Abstract

The mechanisms that control the dynamic composition of RNAPII elongation complexes govern major transitions in the transcription cycle yet are poorly understood. Here, we show that the transcription elongation factor Spt5 determines elongation complex composition to promote productive elongation and the transition to termination. Using an unbiased genetic screen and genomic approaches in *Saccharomyces cerevisiae*, we provide evidence that dephosphorylation of the Spt5 C-terminal repeat domain (CTR) by Glc7/PP1 is required to dislodge the Paf1 complex (Paf1C) from RNAPII near the cleavage and polyadenylation site (CPS). Mutations in Paf1C or the Spt5 CTR that dissociate Paf1C from RNAPII bypass the requirement for two critical regulators of Glc7 in the cleavage and polyadenylation factor that promote Glc7 enrichment at the 3’ ends of genes. Depletion of Glc7 causes aberrant retention of Paf1C past the CPS and a dramatic increase in readthrough transcription, which is fully suppressed by Paf1C mutations. Our results demonstrate that Paf1C retention antagonizes transcription termination and that Glc7-mediated restructuring of the RNAPII elongation complex to evict Paf1C at the CPS is a critical step in the transition from elongation to termination.

---

Many factors associate with RNAPII to coordinate co-transcriptional events and drive major transitions in the transcription cycle. Early in the transition from initiation to elongation, a set of highly conserved transcription elongation factors, including Spt5, Spt6 and Paf1C, assembles onto RNAPII and promotes productive elongation (Aoi and Shilatifard 2023; Farnung 2025). Additional factors engage with this core elongation complex to couple transcription with RNA processing and chromatin regulation and to facilitate the transition from elongation to termination (Aoi and Shilatifard 2023; Rodriguez-Molina et al. 2023; Farnung 2025; Lopez Martinez and Svejstrup 2025). The dynamic association of these factors with RNAPII is often controlled by protein kinases and phosphatases that target the C-terminal domain (CTD) of the largest subunit of RNAPII, Rpb1, or the elongation factor Spt5 (Harlen and Churchman 2017; Sun and Fisher 2025a). For some of these interactions, the temporal controls and structural interfaces have been documented. However, much less is known about the mechanisms that direct changes to the composition of RNAPII elongation complexes or the consequences when these mechanisms fail.

Spt5, in a heterodimeric complex with Spt4 (DSIF in humans) (Wada et al. 1998), is one of the first elongation factors to associate with RNAPII upon promoter escape and travels with RNAPII until termination (Mayer et al. 2010). Spt5 promotes RNAPII processivity, co-transcriptional RNA processing, and chromatin maintenance, and it regulates RNAPII promoter-proximal pausing in metazoans (Sun and Fisher 2025a). Some of these functions are mediated by the Spt5 C-terminal repeat domain (CTR) and its phosphorylation by CDK9 (Bur1 in *S. cerevisiae*) (Yamada et al. 2006; Liu et al. 2009; Zhou et al. 2009; Qiu et al. 2012; Parua et al. 2018; Sun and Fisher 2025a). The CTR is a conserved feature of eukaryotic Spt5, consisting of repeats of a consensus sequence (Swanson et al. 1991; Pei and Shuman 2003; Yamada et al. 2006; Zhou et al. 2009). In *S. cerevisiae*, the CTR contains 15 repeats of S[T/A]WGG[A/Q] in which the first serine residue can be phosphorylated (Swanson et al. 1991; Liu et al. 2009). The CTR serves as an interaction hub for multiple factors, including Paf1C (Liu et al. 2009; Zhou et al. 2009; Mayekar et al. 2013; Wier et al. 2013; Vos et al. 2020; Sun and Fisher 2025b), pre-mRNA capping enzymes (Pei and Shuman 2002; Doamekpor et al. 2014) and cleavage factor (Mayer et al. 2012b).

Paf1C is another highly conserved multi-functional elongation factor associated with RNAPII. In yeast, Paf1C consists of five subunits, Paf1, Ctr9, Cdc73, Rtf1, and Leo1 (Mueller and Jaehning 2002; Squazzo et al. 2002). In metazoans, Paf1C additionally contains Wdr61, and Rtf1 is less tightly associated (Zhu et al. 2005; Vos et al. 2018). In both yeast and mammalian cells, Paf1C directly stimulates transcription elongation, RNAPII processivity, and transcription-coupled histone modifications (Van Oss et al. 2016; Hou et al. 2019; Vos et al. 2020; Zumer et al. 2021; Fetian et al. 2023; Francette and Arndt 2024). The assembly of Paf1C onto RNAPII involves at least three attachment points: Rtf1 interacts with the phosphorylated Spt5 CTR (Liu et al. 2009; Qiu et al. 2012; Mayekar et al. 2013; Wier et al. 2013; Vos et al. 2020), and Cdc73 interacts with both the Rpb1 linker region and Spt6 (Amrich et al. 2012; Ehara et al. 2022; Ellison et al. 2023). Despite directly interacting with Spt5, Spt6, and RNAPII, Paf1C has a unique occupancy pattern on transcribed genes. It joins the elongation complex after Spt5 and Spt6 and dissociates from RNAPII near the CPS, whereas Spt5 and Spt6 remain associated with RNAPII downstream of the CPS in the termination zone (Mayer et al. 2010). The mechanisms that trigger Paf1C dissociation at the CPS and the importance of this step for transcriptional control are unknown.

At the 3’ ends of genes, pre-mRNA cleavage and polyadenylation are coupled to transcription termination. Transcription through the polyadenylation signal slows the rate of RNAPII elongation in parallel with the CTD-mediated recruitment of cleavage and termination factors (Rodriguez-Molina et al. 2023). Upon pre-mRNA cleavage, the 5’ end of the RNAPII-associated transcript is degraded by the 5’-to-3’ exoribonuclease Xrn2 (Rat1 in *S. cerevisiae*), leading to destabilization of the RNAPII elongation complex and transcription termination (Zeng et al. 2024). Spt5 and the phosphorylation state of the CTR play critical roles in these processes. Studies in fission yeast and mammalian cells have shown that Protein Phosphatase 1 (PP1) dephosphorylates the Spt5 CTR at the 3’ ends of genes (Kecman et al. 2018; Parua et al. 2018; Cortazar et al. 2019). In *S. cerevisiae*, the PP1 ortholog, Glc7, is a component of the phosphatase/APT module of the cleavage and polyadenylation factor complex (CPF) (Nedea et al. 2003; Nedea et al. 2008; Casanal et al. 2017). Importantly, dephosphorylation of Spt5 by PP1 slows the rate of elongation and primes RNAPII for Xrn2-mediated termination (Cortazar et al. 2019). However, the mechanism by which Spt5 dephosphorylation translates into a change in elongation rate remains unknown.

Here, we provide evidence that dephosphorylation of the Spt5 CTR by Glc7, the sole PP1 in *S. cerevisiae*, promotes dissociation of the positive elongation factor Paf1C from RNAPII and this is an essential step in transcription termination. Acute Spt5 depletion and mutations that prevent CTR phosphorylation cause widespread loss of Paf1C from chromatin, while a phosphomimetic CTR mutant aberrantly retains Paf1C past the CPS and causes increased transcriptional readthrough. Mutations that uncouple Paf1C from RNAPII bypass the requirement for the Glc7 phosphatase submodule of the CPF. Moreover, depletion of Glc7 from the nucleus prevents the timely dissociation of Paf1C, Spt5 and RNAPII from chromatin and greatly impairs transcription termination. Remarkably, the deletion of Paf1C subunits fully suppresses transcriptional readthrough at both protein-coding and snoRNA genes caused by nuclear depletion of Glc7. We conclude that the essential function of Glc7 in the context of the CPF is to dislodge Paf1C from RNAPII to enable the elongation-to-termination transition.

## Results

### Acute depletion of Spt5 broadly impacts RNAPII elongation complex assembly and transcription

To investigate the direct roles of Spt5 in RNAPII transcription, we used an auxin inducible degron (AID) (Nishimura et al. 2009; Yesbolatova et al. 2020) to rapidly deplete Spt5 followed by chromatin immunoprecipitation sequencing (ChIP-seq) and 4-thiouracil (4tU) labeling of nascent transcripts (4tU-seq) (Miller et al. 2011). The miniAID tag on Spt5 (mAID-Spt5) did not interfere with Spt5 function prior to depletion, as the strains grew normally and did not exhibit an Spt^-^ phenotype (**Supplemental Fig. S1A**). Upon addition of 5-Ph-IAA (Yesbolatova et al. 2020), mAID-Spt5 was rapidly degraded with little to no change in the levels of Rpb1, Spt6 or Paf1C (**Fig. 1A and Supplemental Fig. S1B,C**). We used 45 minutes as the depletion time for all genomic experiments.

**Fig. 1:**
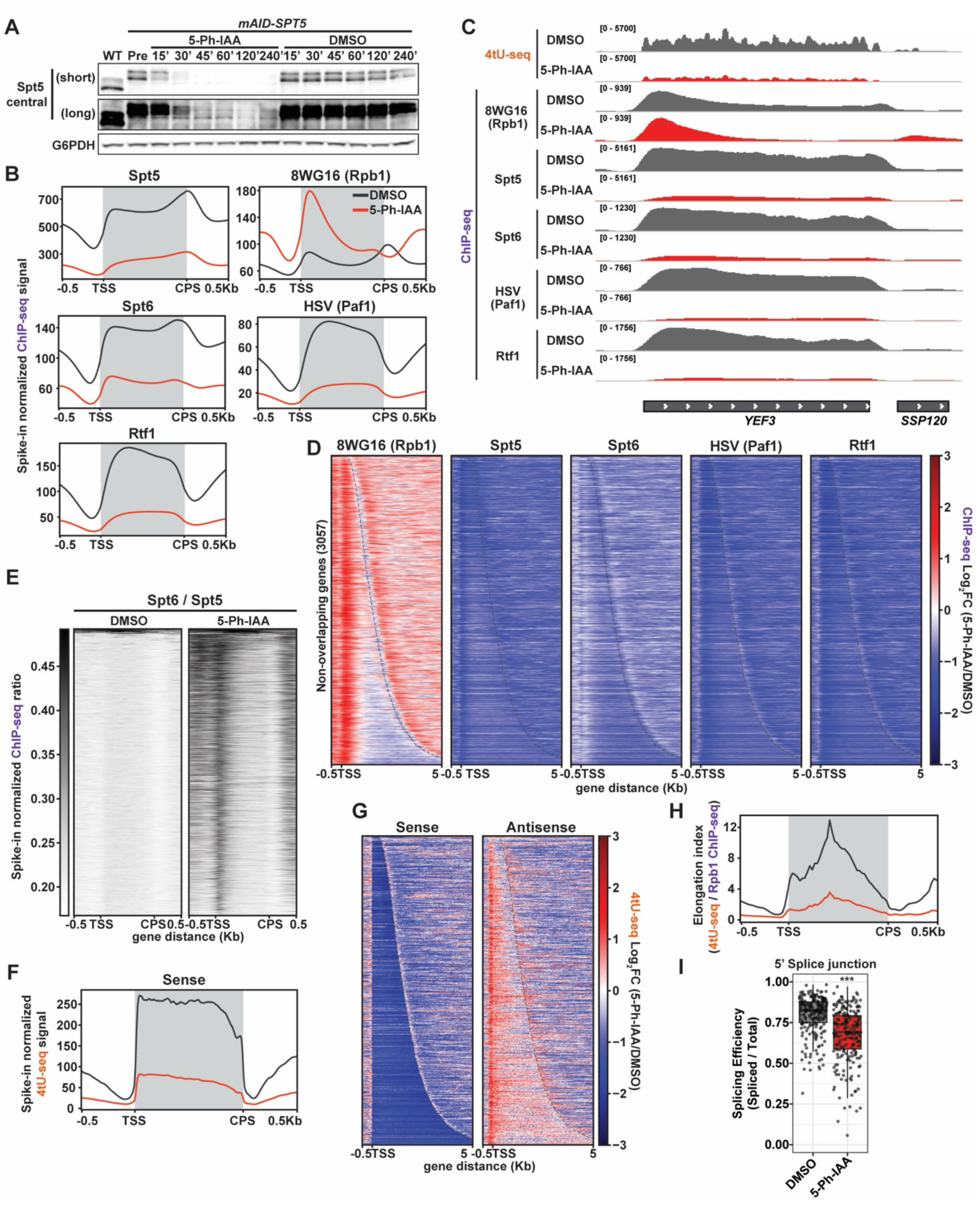
Spt5 is required for RNAPII elongation complex assembly and productive transcription. (A) Western blot analysis of mAID-Spt5 depletion. Controls include a strain lacking the mAID tag on Spt5 (WT), cells treated with vehicle (DMSO), and cells collected before addition of 5-Ph-IAA or DMSO (Pre). G6PDH serves as a loading control. Short and long exposures are shown. n=3 (B) Metaplots of mean spike-in normalized ChIP-seq signal for the indicated proteins over non-overlapping *S. cerevisiae* genes (n=3,057) (Reim et al. 2020) scaled from transcriptional start sites (TSS) to cleavage and polyadenylation sites (CPS). Shading highlights the region between the annotated TSS and CPS. *mAID-SPT5* cells were treated with 5-Ph-IAA (red) or DMSO (black) for 45 min. Color scheme is the same throughout the figure. n=2 (C) 4tU-seq sense-strand (top) and ChIP-seq signal for the indicated antibodies (below) at *YEF3*, a highly transcribed locus. (D) Heatmaps of log2-fold change in ChIP-seq signal for depleted (5-Ph-IAA) vs. undepleted (DMSO) conditions as in (B) with genes sorted by length. Dashed line indicates CPS. (E) Heatmap of Spt6 ChIP-seq signal relative to Spt5 signal. (F and G) Metaplot (F) and differential heatmap (G) of mean spike-in normalized 4tU-seq signal for sense and antisense transcripts for genes shown in (B). n=2 (H) Metaplot of elongation index. (I) Boxplot of splicing efficiency at the 5’ splice junction of introns (n = 231) per Tukey method. Significance from Mann-Whitney U test with FDR correction: ***p < 0.001.

To study the impact of Spt5 on RNAPII elongation complex composition, we measured Rpb1, Paf1C and Spt6 occupancy upon Spt5 depletion by ChIP-seq (replicate correlations in **Supplemental Table 1**). Spt5 depletion caused RNAPII accumulation at the 5’ ends of genes and a reduction in gene bodies, as previously reported (**Fig. 1B-D**) (Shetty et al. 2017; Henriques et al. 2018; Badjatia et al. 2021; An et al. 2024; Kus et al. 2025). The increased Rpb1 signal downstream of the CPS (**Fig. 1B-D**) is due to the 5’ accumulation of RNAPII at neighboring downstream genes, as confirmed by a comparison of tandem and convergent genes (**Supplemental Fig. S1D**). Depletion of Spt5 also led to a dramatic loss of the Paf1C subunits Paf1 and Rtf1 from chromatin (**Fig. 1B-D and Supplemental Fig. S1E**). Consistently, bulk levels of Paf1C-dependent histone modifications, H2BK123ub and H3K4me3, also decreased at longer times of depletion (**Supplemental Fig. S1B**). Furthermore, we observed a strong reduction in Spt6 occupancy, revealing a role for Spt5 in recruiting or retaining Spt6 within the elongation complex (**Fig. 1B-D and Supplemental Fig. S1E**). If Spt6 recruitment is dependent on Spt5, we expect that the Spt6:Spt5 ChIP-seq signals would remain unchanged after Spt5 depletion. However, Spt6 occupancy normalized to Spt5 showed an enrichment at the 5’ ends of genes, similar to the RNAPII profile, suggesting that Spt6 is recruited to RNAPII, to some extent, in the absence of Spt5 (**Fig. 1E**). Given the importance of Spt6 for Paf1C recruitment/retention (Aoi et al. 2022; Ellison et al. 2023), the loss of Paf1C occupancy upon Spt5 depletion could reflect the combined effects of removing both Spt5 and Spt6 from chromatin.

The effect of Spt5 depletion on transcription, assayed by spike-in normalized 4tU-seq, revealed several changes (replicate correlations in **Supplemental Table 2**). First, there was a drastic decrease in sense RNA synthesis as well as an increase in antisense transcription concentrated at the 5’ ends of genes consistent with previous literature (**Fig. 1C,F and G**) (Shetty et al. 2017; An et al. 2024). Calculation of the elongation index as a proxy for elongation rate by dividing the 4tU-seq signal by RNAPII ChIP-seq signal (Caizzi et al. 2021) showed a severe elongation defect (**Fig. 1H**). Second, our 4tU-seq data revealed a widespread splicing defect upon acute loss of Spt5 (**Fig. 1I and Supplemental Fig. S1F**). Intron retention was previously observed at genomic and individual gene levels in Paf1C mutants (Francette and Arndt 2024), upon Spt5 depletion (Shetty et al. 2017; Maudlin and Beggs 2019), and in a recruitment-defective *spt6* mutant (Connell et al. 2022). Considering the crucial role of Spt5 in RNAPII elongation complex assembly, the profound transcriptional defects caused by Spt5 depletion likely arise from the loss of direct Spt5 functions compounded by the effects of also losing Spt6 and Paf1C functions.

### The Spt5 CTR is critical for genome-wide occupancy of Paf1C and proper transcription elongation

Given the severe consequences of Spt5 depletion, we next investigated the contributions of the CTR and its phosphorylation. First, we performed ChIP-seq on a panel of *spt5* CTR mutants (replicate correlations in **Supplemental Table 3**). The *spt5* mutations either delete the CTR (*spt5ΔCTR* and *spt5Δ5*) or alter the site of phosphorylation, replacing the first serine in all 15 repeats with alanine (*spt5-S1-15A*) or, as a phosphomimetic, aspartic acid (*spt5-S1-15D*) (**Fig. 2A**). The *spt5Δ5* mutation, included as a more severe mutation in our genetic screens (below), removes a portion of the KOW5 domain. To different extents, Spt5 occupancy is reduced in the mutants, which for three of them can be at least partially explained by decreased protein levels (**Fig. 2B,C and Supplemental Fig. S2A**). RNAPII occupancy (Rpb1 and Rpb3) is globally elevated in the CTR truncation mutants but not the substitution mutants (**Fig. 2C and Supplemental Fig. S2B**). While bulk levels of Rpb1, as detected with antibodies against the unphosphorylated CTD (8WG16) and Rpb1 core (Y80), are modestly elevated in the *spt5-S1-15A*, *spt5Δ5*, and *spt5ΔCTR* mutants, we observed no significant changes in the levels of Rpb1 phosphorylated at Ser2 or Ser5 of the CTD in any of the *spt5* mutants (**Fig. 2B and Supplemental Fig. S2A**). When normalized to the Rpb3 ChIP-seq signal, all *spt5* mutants have reduced Spt5 occupancy (**Supplemental Fig. S2C**).

**Fig. 2:**
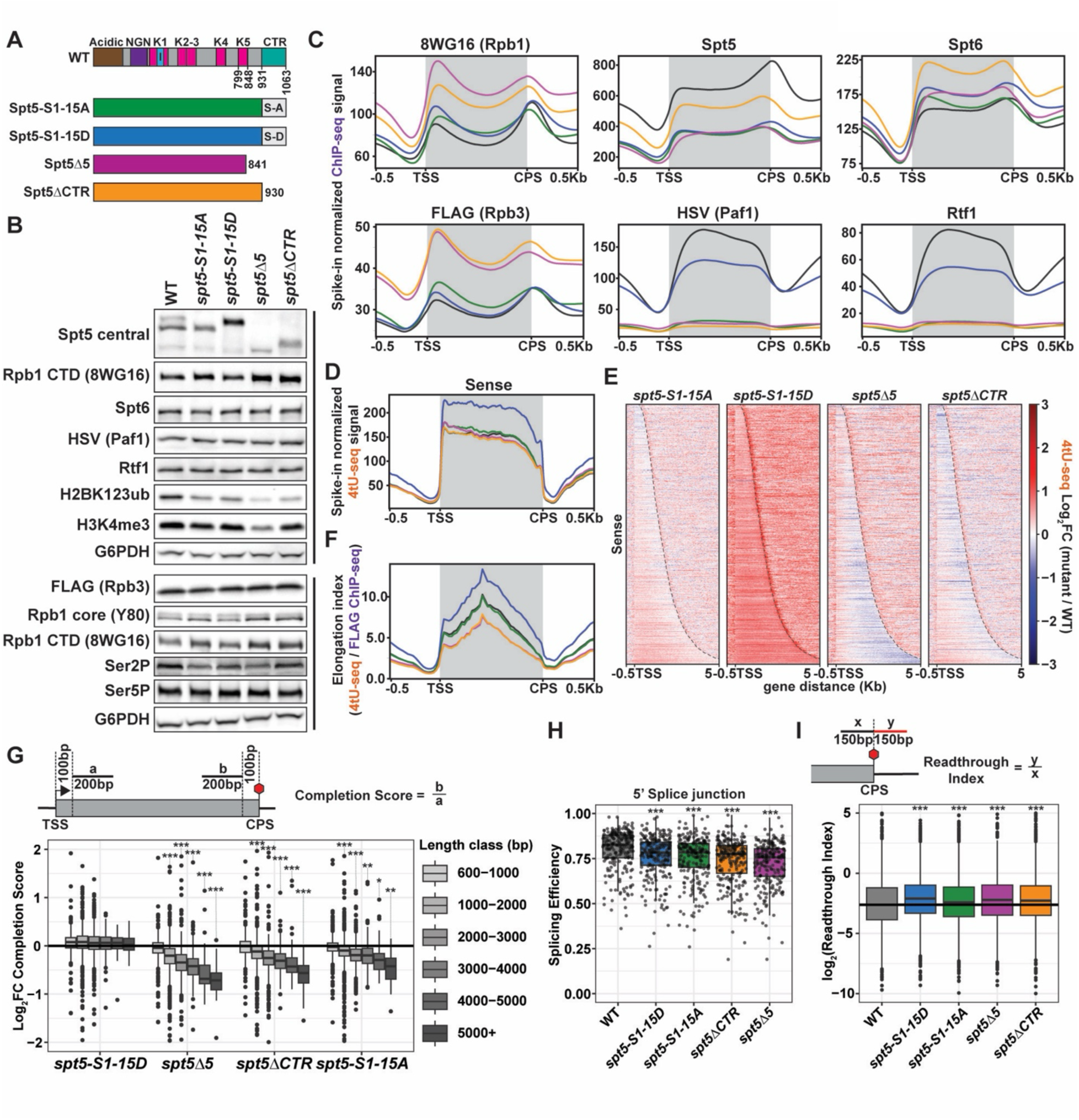
The Spt5 CTR and its phosphorylation regulate multiple aspects of RNAPII transcription. (A) Spt5 mutants used for ChIP-seq and 4tU-seq experiments. Color code is used throughout the figure. Diagram at the top illustrates the domain structure of *S. cerevisiae* Spt5 with domains/regions labeled as NGN=NusG N-terminal homology, I=insertion, and K=KOW.(Meyer et al. 2015) (B) Western blot analysis of *spt5* mutant strains. Black bar indicates the same strains/replicates. n=3 (C and D) Metaplots of mean spike-in normalized ChIP-seq signal for the indicated proteins (C) and 4tU-seq signal for the sense-strand (D) over non-overlapping genes. n=2 (E) Heatmap representation of log2-fold change in 4tU-seq sense-strand signal for *spt5* mutants relative to WT. (F) Metaplot of elongation index. (G-I) Boxplots report the log2-fold change in completion score relative to WT, calculated from 4tU-seq signals as shown in the diagram and stratified by gene-length class (G), distribution of splicing efficiency over the 5’ splice junction of introns (n = 231) (H), and log2 of readthrough index, calculated from 4tU-seq signals as shown in the diagram (I), with outliers shown per Tukey method. Black line in (I) indicates median readthrough index of WT. Significance from Mann-Whitney U test with FDR correction: *p < 0.05, **p < 0.01, and ***p < 0.001.

All non-phosphorylatable *spt5* mutants, including *spt5-S1-15A*, *spt5Δ5* and *spt5ΔCTR*, have greatly reduced levels of Paf1 and Rtf1 on chromatin, even when normalized to account for mutant-specific differences in RNAPII and Spt5 occupancy (**Fig. 2C and Supplemental Fig. S2B-D**). In contrast, the phosphomimetic *spt5-S1-15D* mutant maintains an intermediate level of Paf1 and Rtf1 occupancy on gene bodies and higher levels of Paf1C association per unit of chromatin-bound Spt5 (**Fig. 2C and Supplemental Fig. S2B,D**). These results strongly support a widespread role for CTR phosphorylation in Paf1C recruitment, consistent with structural studies (Wier et al. 2013; Vos et al. 2020). In contrast to Paf1C, Spt6 occupancy levels largely mirror RNAPII occupancy in all the mutants (**Fig. 2C and Supplemental Fig. S2B,C**), indicating that Spt6 occupancy depends primarily on the Spt5 core. Importantly, all *spt5* mutants have bulk levels of Paf1, Rtf1, and Spt6 that are comparable to wild-type levels (**Fig. 2B**). Interestingly, the non-phosphorylatable *spt5* mutants exhibit an upstream shift of the 3’ peaks of RNAPII, Spt5, and Spt6 ChIP-seq signals, suggesting changes in RNAPII elongation properties in this region and potential effects on termination (**Fig. 2C and Supplemental Fig. S2B,E and F**).

Spike-in normalized 4tU-seq analysis of the *spt5* CTR mutants showed distinct effects on transcription (replicate correlations in **Supplemental Table 2**). While the non-phosphorylatable mutants are broadly similar to wild type with respect to global 4tU-seq signal levels, the *spt5-S1-15D* mutant exhibits considerably higher transcription (**Fig. 2D,E**).

Elongation index calculations revealed elongation rate defects only in the truncation mutants (**Fig. 2F**). Conversely, the higher elongation index for the *spt5-S1-15D* mutant indicates a greater frequency of RNAPII passage through the gene. To quantify the impact of CTR perturbations on RNAPII processivity, we calculated a completion score from the 4tU-seq data (Narain et al. 2021; Francette and Arndt 2024). With the notable exception of *spt5-S1-15D*, the *spt5* CTR mutations all cause RNAPII processivity defects (**Fig. 2G**). This phenotype correlates with loss of Paf1C from the elongation complex, as depletion of Paf1C subunits impairs RNAPII processivity (Zumer et al. 2021; Francette and Arndt 2024). Additionally, all mutants exhibit increased levels of unspliced transcripts (**Fig. 2H**). Given the roles of the CTR in pre-mRNA cleavage and RNAPII termination (Mayer et al. 2012b; Baejen et al. 2017; Parua et al. 2018; Cortazar et al. 2019), we calculated a readthrough index. All *spt5* mutants exhibit readthrough transcription to varying degrees (**Fig. 2I and Supplemental Fig. S2G**). Notably, the *spt5-S1-15D* mutant shows the most severe termination defect, accompanied by increased Paf1C retention (**Supplemental Fig. S2G,H**). Together, our analysis of Spt5 acute depletion and *spt5 CTR* mutants underscores the role of Spt5 as a nexus of RNAPII elongation complex assembly to promote elongation, transcript processing, and the transition to termination.

### Genetic screens identify genes that functionally interact with the Spt5 CTR

To interrogate the functions of the CTR and its phosphorylation through the identification of genetic interactors, we performed a high-throughput, transposon-based yeast genetic screen, SAturated Transposon Analysis in Yeast (SATAY) (Michel et al. 2017; Michel and Kornmann 2022). Since most transposon insertions create a loss-of-function allele, higher transposon coverage in a given gene in the mutant background relative to the wild-type background indicates suppression while lower coverage indicates synthetic growth impairment or lethality.

We performed SATAY screens on *spt5Δ5*, *spt5-S1-15A*, and *spt5ΔCTR* mutants. The transposon maps and principal component analysis show high reproducibility between replicates (**Fig. 3A and Supplemental Fig. S3A**).

**Fig. 3:**
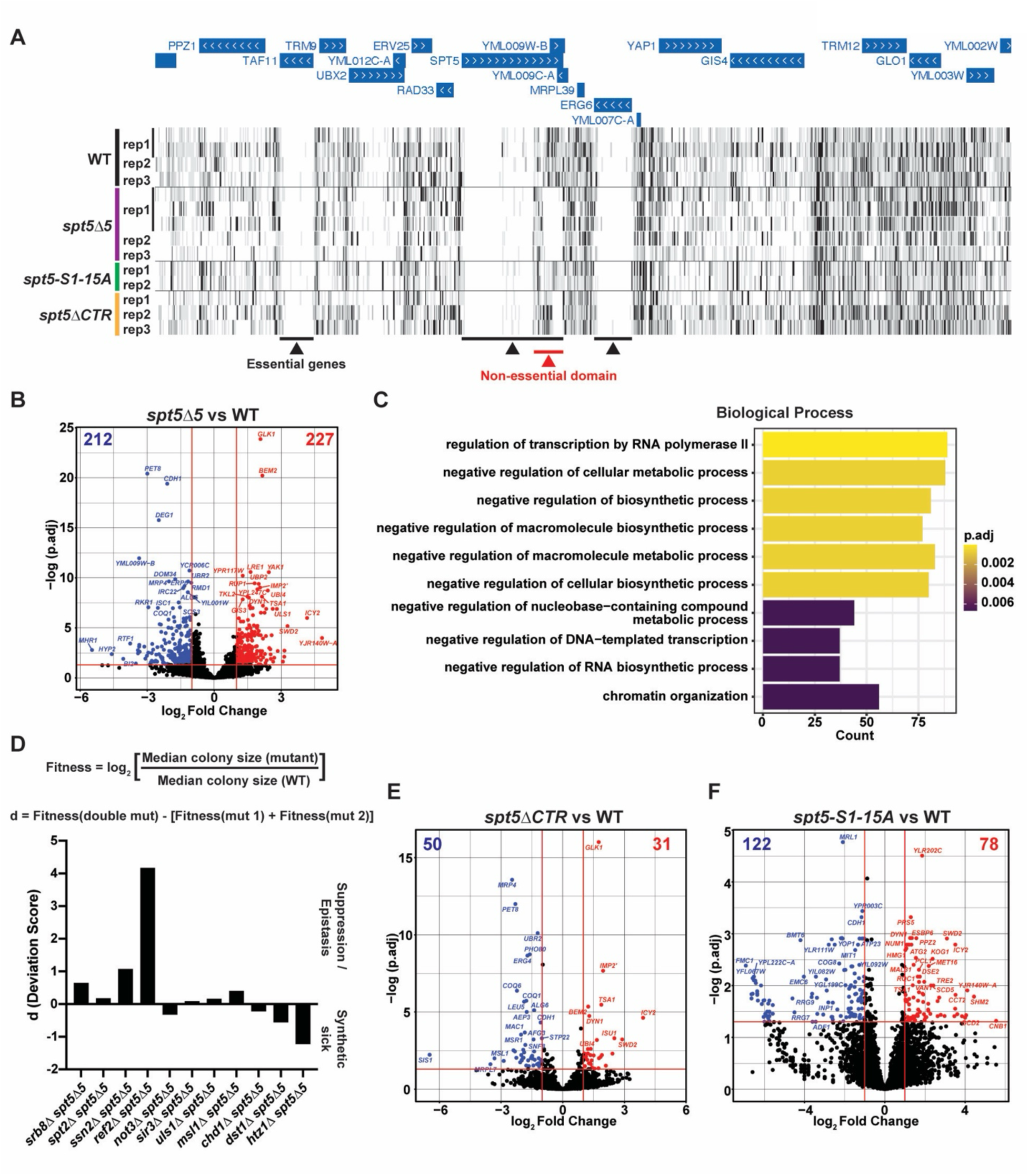
Transposon mutagenesis screens identify genetic interactions with *spt5* CTR mutations. (A) UCSC genome browser tracks showing transposon coverage at *SPT5* and neighboring genes in strains expressing WT or mutant derivatives of Spt5. Rep numbers indicate independent biological replicates. For some biological replicates, multiple technical replicates were tested. (B) Volcano plots showing genes with significantly different numbers of transposon insertions in *spt5Δ5* mutant compared to WT as defined by DESeq2. p.adj < 0.05 and log2FC > 1 or log2FC < -1. To account for differences in transposition efficiency, the number of transposons per gene (tn/gene) was normalized to the total number of transposons in each replicate. (C) Bar graph showing top ten overrepresented Gene Ontology (GO) (Gene Ontology et al. 2023) terms associated with total genes with significantly different tn/gene in the *spt5Δ5* mutant. (D) Bar graph showing deviation score for double mutants of candidate gene deletions with *spt5Δ5*. Fitness was calculated from quantified colony size (in pixels) on yeast tetrad dissection plates. A positive deviation score indicates genetic suppression or epistatic genetic interaction, while a negative deviation score indicates a synthetic impaired growth defect. (E and F) Volcano plots showing transposon insertions in *spt5ΔCTR* (E) or *spt5-S1-15A* (F) mutants and plotted as described in (B).

Focusing first on candidates from the screen with the *spt5Δ5* mutant, which appears especially sensitive to genetic changes, the SATAY results revealed many genes with significant enrichment in transcription- and chromatin-associated gene ontology (GO) terms (**Fig. 3B,C**). We generated deletion mutations *de novo* for 14 genes (**Supplemental Fig. S3B**) to independently test their genetic interactions with the *spt5Δ5* mutation. To this end, we performed growth assays on progeny after yeast tetrad dissection and tested for effects on the Spt^-^ phenotype of *spt5Δ5* (**Supplemental Fig. S3C,D**). We also calculated fitness and deviation scores to evaluate the genetic interactions (**Fig. 3D**) (Duan et al. 2025). For many of these genes, independent testing either by dilution spot assays or quantitation of colony size after tetrad dissection verified the SATAY predictions of positive or negative genetic interactions. In some cases, the candidate mutation or *spt5Δ5* was epistatic to the other mutation (**Fig. 3D and Supplemental Fig. S3D**). Here, we focused on two candidates that exhibited the strongest genetic interactions with *spt5Δ5* and other *spt5* CTR mutations.

Specifically, our SATAY screen uncovered strong positive genetic interactions between *spt5* CTR mutations and mutations targeting the CPF, specifically its phosphatase/APT module (**Supplemental Fig. S4A**). Compared to wild type, we observed a significantly higher number of transposon insertions in the *REF2* gene in the *spt5τι5* background (**Fig. 3D and Supplemental Fig. S4B**) and in the *SWD2* gene in all three *spt5* mutant backgrounds (**Fig. 3B,E,F and 4A**).

*REF2* and *SWD2* encode critical regulatory subunits of Glc7 in the phosphatase/APT module (Nedea et al. 2003; Cheng et al. 2004; Nedea et al. 2008; Casanal et al. 2017; Lidschreiber et al. 2018). Whereas little to no transposition in *REF2* and *SWD2* in the wild type indicates essentiality, increased transposition in the mutants indicates suppression of the severe growth defects associated with the loss of Ref2 and Swd2 by the *spt5* mutations. Overall, the identification of two components of the Swd2-Ref2-Glc7 submodule in our screen points to a specific functional interaction between the Spt5 CTR and the phosphatase/APT module of the CPF.

### Disruption of Paf1C recruitment to RNAPII suppresses loss of Swd2 and Ref2

Building on the SATAY results, we further investigated the genetic suppression of *ref2* and *swd2* mutations by *spt5* mutations. First, we generated *swd2Δ* and *ref2Δ* alleles *de novo* and confirmed their lethality and extreme growth defects, respectively (**Supplemental Fig.**

**S4C**). Next, we generated double mutants of *swd2Δ* or *ref2Δ* with the *spt5* mutations to confirm the genetic interaction results (**Fig. 4B,C**). Consistent with the SATAY results, all non-phosphorylatable *spt5* mutations rescue the lethality/extreme growth defects of the *swd2Δ* and *ref2Δ* mutants. Conversely, the phosphomimetic *spt5-S1-15D* mutation partially rescues the growth defect of the *ref2Δ* mutant but not *swd2Δ* lethality. Our ChIP-seq results showed that Paf1C occupancy is severely reduced in the non-phosphorylatable *spt5* mutants, whereas the *spt5-S1-15D* mutant has an intermediate level of Paf1C occupancy (**Fig. 2C**). Therefore, we tested if Paf1C mutations could phenocopy *spt5* mutations with respect to suppression of *swd2Δ* and *ref2Δ*. Indeed, both *paf1Δ* and *rtf1Δ* strongly suppress the *ref2Δ* and *swd2Δ* mutations (**Fig. 4D**). Given the less severe nature of *ref2Δ* compared to *swd2Δ*, the partial reduction in Paf1C occupancy in the *spt5-S1-15D* mutant might be sufficient to suppress *ref2Δ* but not *swd2Δ*.

**Fig. 4:**
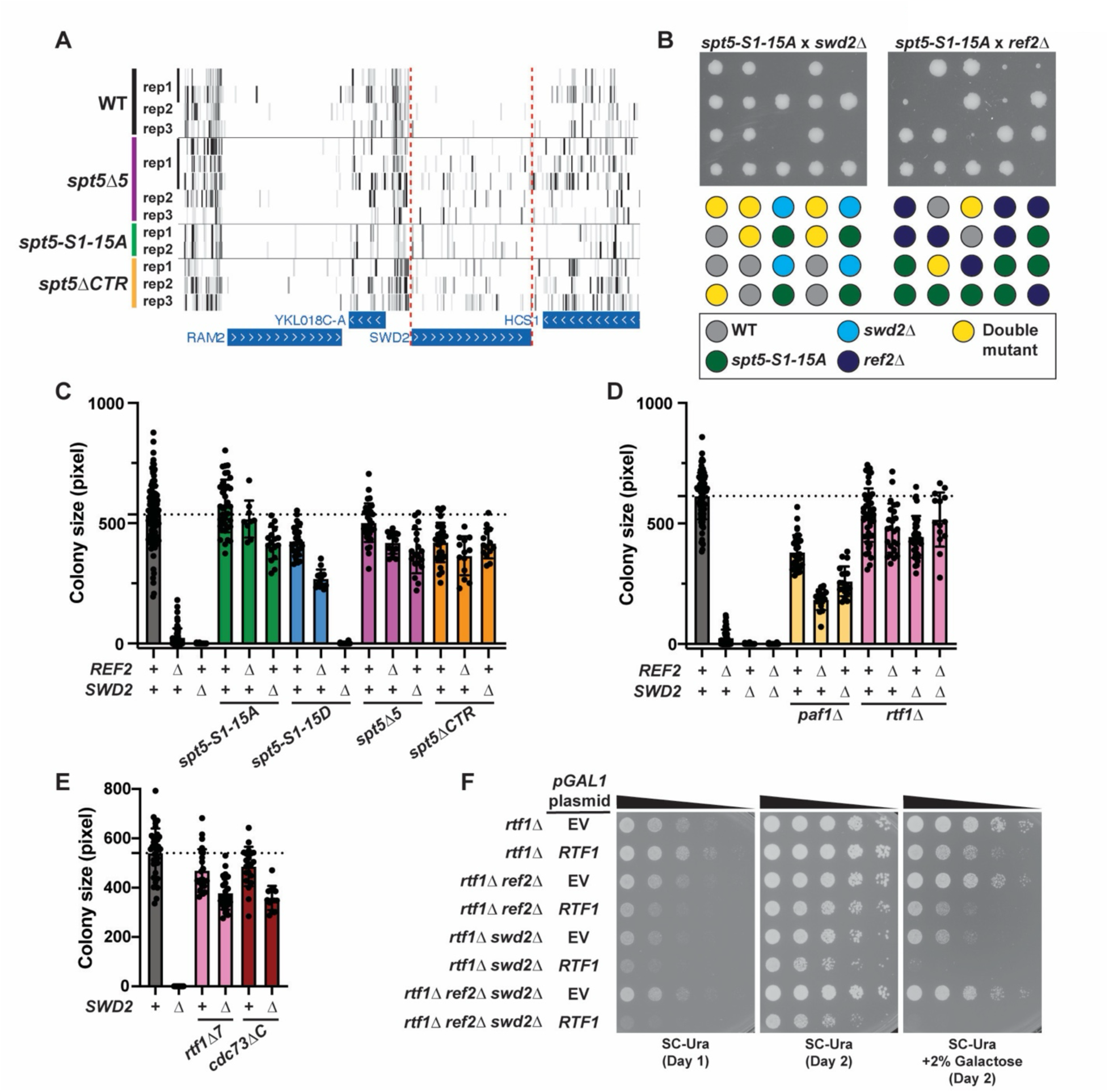
Mutations disrupting Paf1C recruitment to RNAPII rescue the inviability of *swd2Δ* and *ref2Δ* mutants. (A) UCSC genome browser tracks showing transposon coverage in the *SWD2* gene in WT or *spt5* mutant backgrounds. Rep numbers indicate independent biological replicates. For some biological replicates, multiple technical replicates were tested. (B) Tetrad dissection of yeast diploids heterozygous for *spt5-S1-15A* and *ref2Δ* or *swd2Δ*. (C-E) Colony size quantification of yeast tetrad dissections from diploids heterozygous for *ref2Δ* and/or *swd2Δ* with *spt5* mutations (C), Paf1C mutations (D), or Paf1C recruitment-defective mutations (E). WT, *refΔ*, and *swd2Δ* single mutant colonies from all dissections are plotted for each graph. Bars represent mean colony size with standard deviation. Colony sizes were quantified in pixels from scanned images after 3 days of growth on YPD. Black dotted line indicates mean colony size for WT strains. (F) Yeast spot dilution growth assay showing the effect of *RTF1* complementation in the indicated strains. For the plate shown on the right, galactose was added to induce *RTF1* expression from the *GAL1* promoter on the plasmid. EV=empty vector.

Since Paf1C is a multifunctional complex (Francette et al. 2021), we investigated the function of Paf1C that, when defective, overcomes the lethal consequences of deleting *SWD2*. To this end, we tested *rtf1* separation-of-function mutations for *swd2Δ* suppression. These *rtf1* mutations disrupt its interaction with the chromatin remodeler Chd1 (*rtf1-8-9A*) (Tripplehorn et al. 2025), ability to stimulate H2BK123ub (*rtf1-108-110A*) (Van Oss et al. 2016; Fetian et al. 2023), or interaction with the Spt5 CTR for recruitment (*rtf1Δ7*) (Warner et al. 2007). The first two mutations did not have any effect on *swd2Δ* lethality, whereas strong suppression was observed with the *rtf1Δ7* mutation (**Fig. 4E and Supplemental Fig. S4D**). Suppression of *swd2Δ* was also observed for the Paf1C recruitment-defective *cdc73ΔC* mutation, which removes a domain in Cdc73 that interacts with Spt6 and Rpb1 (**Fig. 4E**) (Ellison et al. 2023). Consistent with the tetrad dissection results, conditional expression of *RTF1* from a galactose-inducible plasmid restored *swd2Δ* and *ref2Δ swd2Δ* lethality (**Fig. 4F**). Even low levels of leaky *RTF1* expression from this plasmid in glucose conditions impaired growth in the context of *swd2Δ* and *ref2Δ* mutations. Collectively, these results strongly suggest that, in the absence of a functional Swd2-Ref2-Glc7 submodule, the association of Paf1C with RNAPII is lethal.

### Swd2 and Ref2 are required for Glc7 occupancy at the 3’ ends of genes

Biochemical evidence shows that loss or mutation of Ref2 or Swd2 leads to dissociation of Glc7 from the CPF (Nedea et al. 2003; Nedea et al. 2008; Carminati et al. 2023). Therefore, we hypothesized that Glc7 would fail to associate with chromatin in the absence of Ref2 and Swd2, particularly at the 3’ ends of genes. To test this, we took advantage of the viable *rtf1Δ ref2Δ swd2Δ* triple mutant to profile Glc7, RNAPII, and Spt5 occupancies by ChIP-seq (replicate correlations in **Supplemental Table 4**). Western blot analysis showed comparable levels of RNAPII, Spt5, and HA-Glc7 in the triple mutant and the *rtf1Δ* control strain (**Fig. 5A**). In agreement with Swd2 also being a member of the Set1 histone methyltransferase complex (Cheng et al. 2004), H3K4me1 was undetectable in the triple mutant compared to a highly reduced level in the *rtf1Δ* strain. Our ChIP-seq results revealed Glc7 enrichment at the 3’ ends of genes with peak occupancy just downstream of the CPS in the wild-type and *rtf1Δ* strains and a strong depletion of Glc7 from the 3’ ends of genes in the triple mutant (**Fig. 5B-D**). These observations support a requirement for Swd2 and Ref2 in tethering Glc7 to the CPF *in vivo*.

**Fig. 5:**
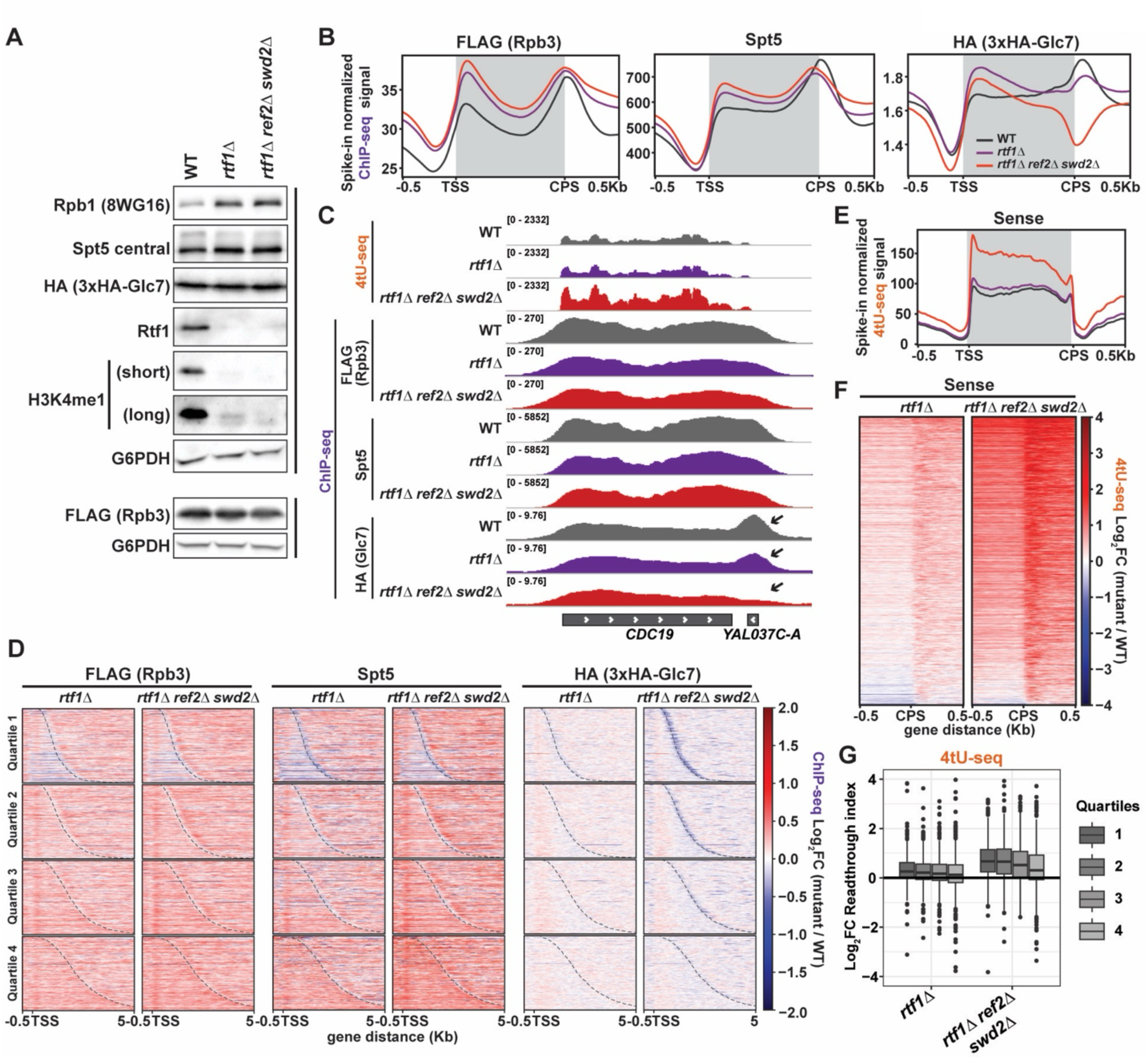
Ref2 and Swd2 are required for chromatin association of Glc7 at the 3’ ends of genes. (A) Western blot analysis of strains used for genomic experiments. Black bar indicates the same strains/replicates. n=3 (B) Metaplots of mean spike-in normalized ChIP-seq data for the indicated proteins. The same color code is used in (C) and (E). n=2 (C) 4tU-seq sense strand and ChIP-seq signal at a highly transcribed locus, *CDC19*. Black arrows indicate Glc7 enrichment peak. (D) Differential heatmaps of ChIP-seq signal for the indicated proteins. Genes are divided into quartiles based on Rpb1 occupancy (highest occupancy in quartile 1) in WT (KY4803) and sorted by gene length. Dashed line indicates CPS. (E) Metaplot showing mean spike-in normalized 4tU-seq signal for the sense strand. Line colors are defined in (B). n=2 (F) Heatmap showing log2-fold change in 4tU-seq signal for the sense strand, centered at the CPS. (G) Boxplot of log2-fold change in readthrough index stratified by quartiles of genic RNAPII occupancy, determined as in (D).

Chromatin occupancy of RNAPII (Rpb3 and Rpb1) and Spt5 was mildly elevated in an *rtf1Δ*-dependent manner, potentially due to slower RNAPII elongation (**Fig. 5B and Supplemental Fig. S5A**). Interestingly, the Spt5 ChIP-seq signal normalized to RNAPII drops off earlier near the CPS in the *rtf1Δ* mutants, suggesting that Spt5 may become prone to dissociation in the absence of Paf1C (**Supplemental Fig. S5B**).

To assess the consequences of Glc7 dissociation from the CPF on transcript synthesis, we performed 4tU-seq on the same set of mutants (replicate correlations in **Supplemental Table 5**). A global increase in sense and antisense transcription was observed in the triple mutant but not the *rtf1Δ* mutant (**Fig. 5C,E, and Supplemental Fig. S5C**). The elongation index points to increased RNAPII passage through gene bodies in the triple mutant (**Supplemental Fig. S5D,E**). Importantly, the triple mutant also showed increased transcriptional readthrough, particularly for the most highly transcribed genes (**Fig. 5F,G**). Defects in splicing and RNAPII processivity were also observed for the triple mutant, expanding the roles of the Swd2-Ref2-Glc7 phosphatase submodule beyond pre-mRNA cleavage and termination (**Supplemental Fig. S5F,G**). Collectively, our results show that Swd2 and Ref2 are required for the localization of Glc7 at the 3’ ends of genes, efficient transcription termination, and suppression of aberrant transcriptional activity within gene bodies.

### Glc7 is required to trigger dissociation of Paf1C from RNAPII at the CPS

Since the Swd2-Ref2-Glc7 phosphatase submodule becomes dispensable for viability in the absence of Paf1C, we hypothesized that the essential function of Glc7 in the CPF is to dislodge Paf1C from RNAPII to facilitate the elongation-to-termination transition. To test this hypothesis, we conditionally depleted Glc7 from the nucleus using the anchor-away system (Haruki et al. 2008) and profiled RNAPII, Spt5 and Paf1C occupancies by ChIP-seq (replicate correlations in **Supplemental Table 6**). Based on a time-course (**Supplemental Fig. S6A**) and the literature (Schreieck et al. 2014), we treated experimental (*GLC7-3xHA-FRB*) and control (*GLC7-3xHA*) strains with rapamycin for one hour.

Glc7 nuclear depletion had no adverse effects on total abundance of RNAPII and elongation factors (**Fig. 6A**). Consistent with work in *S. pombe* and human cells showing PP1-dependent dephosphorylation of Spt5 (Kecman et al. 2018; Parua et al. 2018; Cortazar et al. 2019), Glc7 depletion resulted in Spt5 CTR hyperphosphorylation (**Fig. 6A**). However, for reasons unknown, we also observed a moderate elevation in CTR phosphorylation for the *GLC7-3xHA* control strain and, therefore, included this rapamycin-treated control in all our genomic experiments. Spt5 hyperphosphorylation was also observed in the *glc7-T152K* mutant (Tu and Carlson 1994), further supporting Glc7 as the evolutionarily conserved CTR phosphatase in *S. cerevisiae* (**Supplemental Fig. S6B**).

**Fig. 6:**
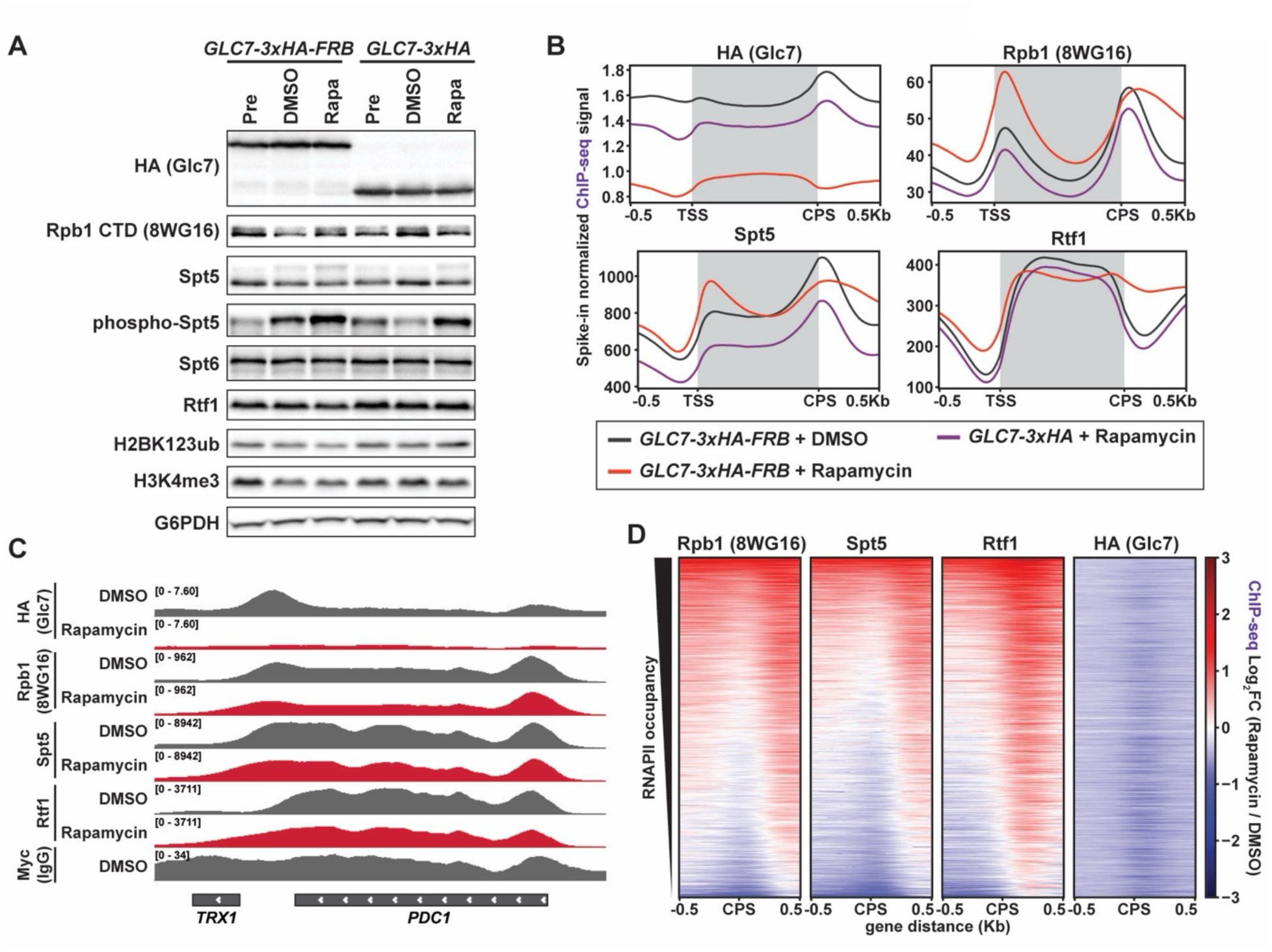
Glc7 is required for the dissociation of Paf1C from RNAPII at the CPS. (A) Western blot analysis showing abundance of indicated proteins and histone modifications in the anchor-away strain (*GLC7-3xHA-FRB*) and a control strain (*GLC7-3xHA*). Cells were untreated (Pre) or treated with DMSO or rapamycin (Rapa, 1µg/mL) for 1 hr. n≥3 (B) Metaplots of mean spike-in normalized ChIP-seq signal for the indicated proteins. n=2 (C) ChIP-seq signals for the indicated proteins at a highly transcribed locus, *PDC1*. Anti-Myc serves as an unrelated antibody control. (D) Differential heatmaps of ChIP-seq data from the *GLC7-3xHA-FRB* strain treated with rapamycin or DMSO and centered at the CPS.

For the *GLC7-3xHA-FRB* strain, Glc7 chromatin occupancy was greatly reduced upon rapamycin treatment as expected (**Fig. 6B-D and Supplemental Fig. S6C,D**). Consistent with previous reports in mammalian cells (Cortazar et al. 2019), *S. pombe* (Kecman et al. 2018; Parua et al. 2018), and *S. cerevisiae* (Schreieck et al. 2014; Collin et al. 2019), we observed 3’ retention of RNAPII upon Glc7/PP1 depletion (**Fig. 6B-D and Supplemental Fig. S6C,D**).

Strikingly, the occupancy levels of Spt5 and Rtf1 downstream of the CPS were also elevated. When normalized to RNAPII and Spt5 ChIP-seq signals, the highest increase in chromatin occupancy downstream of the CPS was observed for Paf1C (**Supplemental Fig. S6E,F**). Importantly, the control strain (*GLC7-3xHA*) for rapamycin treatment and the vehicle-treated (DMSO) Glc7 anchor-away strain showed similar occupancy profiles, confirming that the observed effects are due to nuclear depletion of Glc7 (**Fig. 6B**). Overall, the ChIP-seq results strongly suggest that Glc7 function in the CPF is required for the dissociation of Paf1C, Spt5, and RNAPII during the elongation-to-termination transition.

### Paf1C antagonizes transcription termination

Hypothesizing that nuclear depletion of Glc7 would phenocopy the effects of *swd2Δ* and *ref2Δ* mutations, we asked if *rtf1Δ* and *paf1Δ* mutations could suppress the effects of Glc7 depletion at the level of transcription as measured by 4tU-seq (replicate correlations in **Supplemental Table 7**). Notably, neither *rtf1Δ* nor *paf1Δ* could rescue the lethality caused by nuclear depletion of Glc7, suggesting that Glc7 likely has other essential, CPF-independent functions in the nucleus (**Supplemental Fig. S7A**). Western analysis showed no rapamycin-dependent changes in Ser2P and Ser5P levels, even in the *paf1Δ* strain with highly reduced Ser2P levels, in agreement with previous studies on PP1 (**Supplemental Fig. S7B**) (Schreieck et al. 2014; Kecman et al. 2018). Meanwhile, in each strain, the Spt5 phosphorylation signal consistently increased upon rapamycin treatment (**Supplemental Fig. S7B**).

Nuclear depletion of Glc7 increased both sense and antisense transcription across gene bodies in the wild type (**Fig. 7A and Supplemental Fig. S7C**). In contrast, a global reduction or no change in 4tU-seq signal was observed in the *paf1Δ* and *rtf1Δ* mutants. Consistent with the ChIP-seq data, Glc7 depletion resulted in widespread transcriptional readthrough (**Fig. 7B-D and Supplemental Fig. S7D,E**). The severity of readthrough was positively correlated with gene expression level (**Fig. 7D**). Remarkably, the transcription termination defects brought on by Glc7 anchor-away were almost entirely suppressed by *paf1Δ* and *rtf1Δ* mutations (**Fig. 7B-D and Supplemental Fig. S7D,E**). Given the requirement for Glc7 in snoRNA termination (Nedea et al. 2003; Nedea et al. 2008; Lidschreiber et al. 2018), we also looked at snoRNA gene transcription and found extreme 3’ readthrough upon Glc7 depletion, which was also fully corrected by *paf1Δ* or *rtf1Δ* mutation (**Supplemental Fig. S7F,G**).

**Fig. 7:**
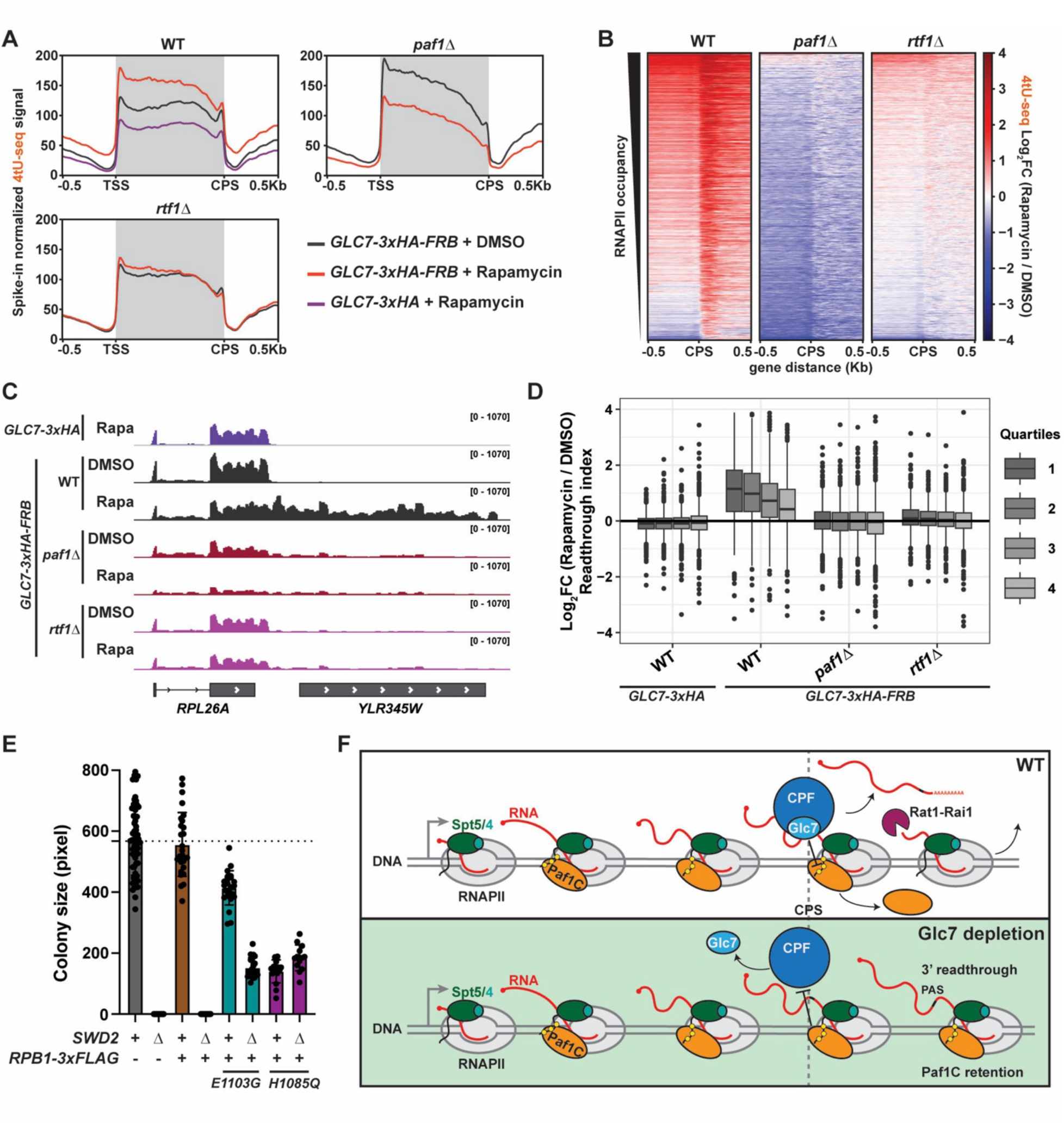
Paf1C mutations strongly suppress transcriptional readthrough caused by nuclear depletion of Glc7. (A) Metaplots of mean spike-in normalized 4tU-seq signal for the sense strand. Cells were treated with DMSO or rapamycin (1µg/mL) for 1 hr. n≥2 (B) Heatmap showing log2-fold change in 4tU-seq signal for the sense strand centered at the CPS upon Glc7 anchor-away. (C) 4tU-seq sense signal at a highly transcribed locus, *RPL26A*. (D) Boxplot showing log2-fold change in readthrough index stratified by genic RNAPII occupancy in quartiles as in Fig. 5G. (E) Colony size quantification of yeast tetrad dissection. Diploids heterozygous for *swd2Δ* and *rpb1* mutations that affect RNAPII catalytic rate were sporulated and haploid progeny were imaged for quantification after 3 days of growth on YPD. (F) Model of the Glc7-mediated elongation-termination transition. Dephosphorylation of the Spt5 CTR (and possibly other substrates) by Glc7 triggers dissociation of Paf1C from RNAPII and proper mRNA cleavage and transcription termination by Rat1-Rai1 exonuclease under WT conditions. Upon Glc7 nuclear depletion, the Spt5 CTR is hyperphosphorylated, leading to Paf1C retention beyond the CPS and transcriptional readthrough. Yellow circles represent phosphates on the Spt5 CTR.

Together, our results demonstrate that inappropriate retention of Paf1C antagonizes transcription termination. As Paf1C is a positive regulator of transcription elongation (Hou et al. 2019; Vos et al. 2020; Zumer et al. 2021; Francette and Arndt 2024), aberrantly fast Paf1C-containing elongation complexes in the termination zone might interfere with the kinetic coupling between transcription elongation and termination (Geisberg et al. 2020). In support of this idea, the *rpb1-E1103G* and *rpb1-H1085Q* mutations, which slow elongation *in vivo* (Malik et al. 2017), can also rescue *swd2Δ* lethality (**Fig. 7E**). These results are consistent with a kinetic competition model (Fong et al. 2015) between transcription-coupled processes governed by Paf1C; however, the dynamics of factor exchange on RNAPII may also contribute to the anti-termination effects of Paf1C retention.

## Discussion

In this study, we provide evidence for a previously unknown role for Glc7 in Paf1C dissociation from RNAPII and demonstrate that this is a necessary step for transcription termination. Through genetic and genomic approaches, we found that Spt5 CTR phosphorylation controls Paf1C residence in the RNAPII elongation complex which, in turn, determines the requirement for Glc7 phosphatase in the CPF. Mutations detaching Paf1C from RNAPII bypass the requirement for the Glc7 submodule in the CPF as well as Glc7 localization at the 3’ ends of genes. Moreover, conditions leading to inappropriate 3’ retention of Paf1C antagonize termination. We conclude that transcription termination requires dissociation of the positive elongation factor Paf1C from RNAPII through Glc7-mediated dephosphorylation of the Spt5 CTR.

Our Spt5 acute depletion experiments demonstrated a global requirement for Spt5 in the chromatin association of Paf1C as well as Spt6 for proper RNAPII elongation complex assembly and transcript synthesis. In the absence of Spt5, Paf1C occupancy is lost across the entire gene body, while Spt6 can still be recruited to the 5’ ends of genes possibly through an interaction with the RNAPII linker (Sdano et al. 2017). Subsequent to recruitment, Spt6 occupancy is not maintained in the absence of Spt5, consistent with Spt5 constituting a large portion of the Spt6 binding surface in the RNAPII elongation complex (Vos et al. 2020; Ehara et al. 2022).

Therefore, the increased antisense transcription detected upon Spt5 depletion likely arises from chromatin dysregulation caused by the combined loss of Spt5 histone binding activity (Evrin et al. 2022) and the histone chaperone activity of Spt6 (Miller et al. 2023).

Our comparative analysis of *spt5* mutations revealed multi-layered regulation of gene expression by the CTR and its phosphorylation, including effects on Paf1C occupancy and RNAPII processivity as well as splicing and termination efficiency. For the truncation mutants, our observations are consistent with previous findings linking the CTR to termination efficiency (Mayer et al. 2012b; Baejen et al. 2017). With respect to the substitution mutants, the *spt5-S1-15D* mutant exhibited more severe transcriptional readthrough than the *spt5-S1-15A* mutant. We propose that this difference in readthrough propensity largely reflects a difference in Paf1C recruitment/retention in these mutants. Indeed, genetic interactors identified in our SATAY screens with *spt5* mutants also genetically interact with Paf1C mutations, highlighting the close functional relationship between Spt5 CTR phosphorylation and Paf1C. Similarly, the *spt5-S1-15D* mutant was uniquely unaffected for RNAPII processivity, suggesting that the Spt5 CTR promotes RNAPII processivity, in large part, through phosphorylation-mediated Paf1C recruitment (Hou et al. 2019; Zumer et al. 2021; Francette and Arndt 2024). However, since the truncation mutants exhibited more severe elongation and mRNA processing defects than the *spt5-S1-15A* mutant despite similar reductions in Spt5 and Paf1C occupancies, our data also suggest Paf1C-independent functions of the CTR in transcription.

The most striking result from our SATAY screen and subsequent genetic experiments is the strong suppression of *swd2* and *ref2* mutations by the absence of Spt5 CTR phosphorylation or by mutations that impair Paf1C recruitment to RNAPII. Mutations that disrupt the interaction between Ref2 and Glc7 are lethal due to the loss of Glc7 from the CPF, emphasizing the essential function of Glc7 in the CPF (Nedea et al. 2008; Carminati et al. 2023). The robust growth of the *rtf1Δ ref2Δ swd2Δ* mutant and the loss of 3’-end enrichment of Glc7 in this strain indicate that the requirement for Glc7 in the CPF can be bypassed by disabling Paf1C. Indeed, nuclear depletion of Glc7 caused retention of Paf1C beyond its normal dissociation site near the CPS and widespread termination defects that were fully reversed by deletion of *PAF1* or *RTF1*. While a termination requirement for PP1 has been reported in mammalian cells (Cortazar et al. 2019), fission yeast (Kecman et al. 2018; Parua et al. 2018) and budding yeast (Schreieck et al. 2014), our results argue that the essential function of PP1 in the CPF is to dislodge Paf1C and its associated anti-termination effects from RNAPII.

Glc7 is also present in the APT complex, which is critical for snoRNA termination in budding yeast (Nedea et al. 2003; Nedea et al. 2008; Lidschreiber et al. 2018). We found that Paf1C mutations also suppress readthrough transcription of snoRNA genes caused by nuclear depletion of Glc7. This is in contrast with reports that Paf1C and its dependent histone modifications are required for efficient snoRNA 3’-end formation (Sheldon et al. 2005; Terzi et al. 2011; Tomson et al. 2011; Tomson et al. 2013; Ellison et al. 2019). In those earlier studies, steady-state transcript levels were measured whereas we used nascent transcriptomics, suggesting that deletion of Paf1C might stabilize snoRNA readthrough transcripts. Interestingly, we also observed a global increase in transcriptional output across gene bodies in the *rtf1Δ ref2Δ swd2Δ* mutant and upon nuclear depletion of Glc7, suggesting that the Swd2-Ref2-Glc7 submodule may negatively regulate transcription elongation. Considering that Paf1C mutations suppress this global increase in transcription in Glc7 depletion conditions, Glc7 may normally down-regulate transcription by limiting the levels of Paf1C associated with RNAPII on gene bodies.

Taken together, our data strongly suggest that aberrant Paf1C retention on RNAPII through Spt5 CTR hyperphosphorylation underlies termination defects caused by loss of Glc7 from the CPF. This would also explain elevated 3’ readthrough in the phosphomimetic *spt5-S1-15D* mutant. We note, however, that the *spt5-S1-15D* mutant does not fully phenocopy nuclear depletion of Glc7 at the level of transcription, possibly due to globally reduced Spt5 occupancy and/or multiple targets of Glc7 in the RNAPII complex including the Rpb1 CTD (Mayer et al.

2012a; Schreieck et al. 2014; Collin et al. 2019). It is also notable that transcription readthrough caused by nuclear depletion of Glc7 was entirely suppressed by *rtf1Δ* and *paf1Δ*, whereas a moderate level of 3’ readthrough was evident in the *rtf1Δ ref2Δ swd2Δ* triple mutant (**Supplemental Fig. S7D-G**). This discrepancy could arise from at least two sources. The triple mutant likely represents an equilibrated state of the viable, suppressed condition and might have less 3’ readthrough than a *ref2Δ swd2Δ* double mutant, an inviable state. Alternatively, Ref2 and Swd2 might have Glc7-independent functions in the CPF. For example, Ref2 is critical for the interaction between APT and RNAPII (Carminati et al. 2023) and therefore CPF recruitment might be compromised in the triple mutant.

At least two non-mutually exclusive mechanisms can explain how Paf1C antagonizes mRNA cleavage and/or transcription termination. (1) RNAPII fails to decelerate past the CPS due to the elongation-promoting activity of Paf1C. In fact, Spt5 dephosphorylation by PP1 in mammalian cells is important for RNAPII deceleration (Parua et al. 2018; Cortazar et al. 2019), and downstream poly(A) sites are favored by fast RNAPII mutants (Geisberg et al. 2020).

Conversely, depletion of Paf1C subunits in mammalian cells leads to a 5’-shift in poly(A) site selection (Yang et al. 2016). (2) Paf1C physically occludes the binding surface on RNAPII for cleavage and termination factors. There is evidence of a direct interaction between RNAPII and CPF (Carminati et al. 2023) and Xrn2 (Zeng et al. 2024; Kus et al. 2025). Although we provide genetic evidence in support of the kinetic competition mechanism, further structural and biochemical studies are needed to test the second mechanism.

Finally, in the context of current literature, we propose a model (**Fig. 7F**) wherein the CPF is recruited by its interaction with Ser2P of the RNAPII CTD (Ahn et al. 2004) at the 3’ ends of genes and triggers dissociation of Paf1C through dephosphorylation of the Spt5 CTR (Kecman et al. 2018; Parua et al. 2018; Cortazar et al. 2019) and likely additional targets such as Tyr1P of the RNAPII CTD (Mayer et al. 2012a; Schreieck et al. 2014; Collin et al. 2019).

Removal of Paf1C slows RNAPII elongation and enables CPF access to RNA and/or RNAPII for efficient pre-mRNA cleavage and transcription termination. Paf1C has been reported to interact with the Cft1 subunit of the CPF (Nordick et al. 2008) and therefore may play a role in recruiting CPF prior to its dissociation. In summary, we have identified the Glc7-mediated dissociation of Paf1C from RNAPII as a pivotal event in transcription termination.

### Limitations of the study

Our work focused on Spt5 CTR dephosphorylation by the Glc7 phosphatase. As the only PP1 in budding yeast, Glc7 has many substrates including other targets in the RNAPII elongation complex, including Tyr1P of the CTD (Schreieck et al. 2014). While our collective data support Spt5 CTR dephosphorylation as a key driver of Paf1C dislodgement, they do not exclude the possibility that dephosphorylation of additional Glc7 substrates in the RNAPII elongation complex contributes to the dissociation of Paf1C at the CPS. Genetic mutants, including the *rtf1Δ ref2Δ swd2Δ* triple mutant and the *spt5* CTR mutants, represent equilibrated states. Therefore, the transcriptional properties of these mutants likely reflect both the direct and indirect effects of the mutations with the latter including cellular compensation mechanisms.

## Materials and methods

### Yeast strains, growth conditions, spot growth assays, and colony size quantification

*S. cerevisiae* strains used in this study (**Supplemental Table 8**) are isogenic to FY2, a *GAL2*^+^ derivative of S288C (Winston et al. 1995). Mutant strains were generated using one-step integrative transformation or derived from genetic crosses. Yeast transformations were performed following the lithium acetate protocol (Rose et al. 1990). Tagging of *GLC7* with *3xHA* or *3xHA-FRB* at either the C- or N-terminus was done using CRISPR/Cas9 as described previously (Laughery et al. 2015). Mutations were confirmed by PCR and/or Sanger sequencing. Sequences of oligonucleotides used for strain constructions are available upon request. Plasmids are listed in **Supplemental Table 9**.

*S. cerevisiae*, *K. lactis,* and *S. pombe* strains were grown at 30°C in YPD medium supplemented with 400 µM tryptophan (YPD+W) or synthetic complete (SC) dropout medium (Rose et al. 1990). For auxin-induced depletion of mAID-Spt5, yeast cultures at an OD600 of ∼0.5 (∼10^7^ cells/mL) were treated with 5-phenyl-indole-3-acetic acid (5-Ph-IAA, Tocris Bioscience, 7392) at a final concentration of 5 µM or an equivalent amount of vehicle (DMSO) and incubated at 30°C for the indicated time. For Glc7 anchor-away experiments, 1 µg/mL rapamycin (MedChemExpress, HY-10219) or an equivalent amount of DMSO was added to the yeast cultures. All cultures harvested for western blots were quenched with cold methanol (stored at -20°C) at ∼33% v/v final concentration to stop treatment at the indicated timepoint and preserve post-translational modifications (Alexander et al. 2010). Cells were washed once with water before centrifugation, and cell pellets were flash frozen in liquid nitrogen.

Serial dilution spot growth assays were performed as described (Francette and Arndt 2024). Plates were incubated at 30°C and scanned (Epson Perfection V600 photo) at ∼24-hour intervals. The colony sizes on yeast tetrad dissection plates were quantified in pixels using a custom R script that utilizes EBImage R package (Pau et al. 2010). Fitness and deviation scores were calculated using the quantified colony sizes as described previously (Duan et al. 2025).

### Western blot analysis

Protein extracts were prepared from methanol-quenched cells (see above) equivalent to an OD600 of 10 using the post-alkaline method (Kushnirov 2000) with modifications. Briefly, cell pellets were resuspended in 100 µL of water or 2x PhosSTOP phosphatase inhibitor stock (Roche, 04906845001) and treated with 100 µL of 0.2 M NaOH for 5 min at room temperature before centrifugation at 14,000 rpm for 5 min. The resulting pellets were boiled in 200 µL of 1x SDS-PAGE loading buffer for 5 min, and centrifuged at 14,000 rpm for 5 min. The proteins in the supernatant were resolved by SDS PAGE for western blotting. Additional experimental details and antibody information are provided in the Supplemental Material.

### ChIP-seq sample preparation and library construction

ChIP was performed with 250 mL cultures of yeast cells grown to an OD600 of 1.0 (∼2 x 10^7^ cells/mL) as described (Ellison et al. 2023). When yeast cultures were exposed to treatments (*e.g.* auxin), formaldehyde was added at the indicated timepoint post treatment. For all ChIP experiments, *S. cerevisiae* chromatin was mixed with spike-in chromatin at a 1:10 ratio (spike-in chromatin : *S. cerevisiae* chromatin) based on protein concentration (Pierce BCA protein assay kit; Thermo Fisher Scientific, 23227). Different sources of spike-in chromatin were used for different IP reactions depending on the antibody cross-reactivity, which was confirmed by western blotting. *S. pombe* KP10 chromatin was used for 8WG16 (BioLegend, 664906, 6 µL/1350 µg) and HSV (Sigma-Aldrich, H6030, 5 µL/1350 µg) IPs in Fig. 1 and 2, while *S. pombe* KP06 chromatin was used for 8WG16 (6 µL/1350 µg or 4 µL/900 µg), HA (Santa-Cruz, sc-7382 AC, 60 µL/1350 µg or 60 µL/1800 µg) and Myc (non-specific IgG control, gift from John Woolford, 5 µL/900 µg) IPs in Fig. 5 and 6. *Kluyveromyces lactis* KL01 chromatin was added to IP reactions for Spt5 central (gift from Grant Hartzog, 2 µL/900 µg or 3 µL/1350 µg), Spt6 (gift from Tim Formosa, (McCullough et al. 2015), 8 µL/1350 µg or 6 µL/900 µg) and Rtf1 ((Squazzo et al. 2002), 4 µL/1350 µg or 3 µL/900 µg), while *K. lactis* KL02 chromatin was used for Rpb3-FLAG (Millipore, A2220, 100 µL slurry/900 µg) IPs. IP reactions were incubated at 4°C overnight on an end-over-end roller. Pre-equilibrated 50% slurry of protein A or G beads (Cytiva, 17-5280-01 or 17-0618-01, 80 µL/1350 µg or 60 µL/900 µg) was added, and samples were incubated at room temperature for 1 hour with rotation. Beads were washed, and IP DNA was eluted and treated with RNase as described (Ellison et al. 2023). DNA was purified using the QIAquick PCR Purification Kit (Qiagen, 28106). ChIP-seq libraries were prepared using the NEBNext Ultra II DNA kit (NEB, E7645S/S) and multiplex oligos (NEB, E6440S/L). AMPure XP beads (Beckman Coulter, A63881) were used for DNA library clean-up. Libraries were paired-end sequenced (2x50 bp) on the Illumina NextSeq 2000 platform.

### 4tU-seq sample preparation and library construction

The protocols for 4tU labeling and cell harvesting were adapted from (Barrass and Beggs 2019; Francette and Arndt 2024). First, strain/mutant-specific linear regression equations of OD600 versus cell number were determined by counting cells under a microscope with a hemocytometer. At the time of the harvest, the number of cells was extrapolated from a specific OD600 using the linear equation. For all cell harvests, a culture-volume equivalent of 5 x 10^8^ or 6 x 10^8^ total cells was transferred to a fresh tube for 4tU labeling. Cells were labelled with 4tU (Sigma-Aldrich, 440736-1G) at a 5 mM final concentration for 5 min at room temperature with shaking and quenched with cold methanol (stored at -20°C) at 33% v/v final concentration to halt 4tU labeling. For mutant and WT strains, cultures were grown to an OD600 of ∼1.0 before 4tU labeling. For strains with treatment, cultures were grown to an OD600 of ∼0.7 before being split and treated, and 4tU was added at the indicated timepoint post treatment for 5 min labeling. Spike-in strain *S. pombe* (KP10) was grown to an OD600 of 1.0 and labelled with 4tU for 10 min. Methanol-quenched cultures were centrifuged at 3000 rpm for 3 min at 4°C, and cell pellets were washed once with ice cold water before being flash frozen in liquid nitrogen. For all cultures, cells were also counted under a microscope using a hemocytometer to record the cell concentration at the time of harvest, which was used for post-hoc cell count normalization.

*S. cerevisiae* cells were mixed with spike-in *S. pombe* cells at a 10:1 ratio (*S. cerevisiae*:*S. pombe*) by cell number. For Glc7 anchor-away samples including two replicates of WT, *paf1Δ* or *rtf1Δ* strains, a *S. cerevisiae* cell suspension volume equivalent to 5 x 10^8^ cells based on cell count at the time of harvest was used as the input. For all other strains including another two replicates of WT Glc7 anchor-away, the entire harvested cell pellet (equivalent to 6 x 10^8^ cells) was used as the input for RNA extraction. Total RNA extraction, followed by biotinylation and isolation of 4tU-labeled RNA, was performed as described (Barrass and Beggs 2019; Francette and Arndt 2024). 4tU-seq libraries were prepared using a Universal Plus Total RNA-seq with NuQuant kit (Tecan, 0361-A01), UPlus Total RNA AnyDeplete module (Tecan, 0370-A01) and UPlus Total AnyDeplete Custom Probe Mix for *S. cerevisiae* (Tecan, S02714). Libraries were sequenced (paired-end, 100 bp) on the Illumina NextSeq 2000 platform.

### SATAY transposon library preparation

Transposon libraries of *SPT5 ade2Δ* and *spt5 ade2Δ* strains were generated and sequenced as described (Michel et al. 2017) (https://satayusers.kornmann.group/complete-protocol). The strains were transformed with pBK549 plasmid (Michel and Kornmann 2022), and individual transformants are considered biological replicates. Each transposon library was generated on ∼50-60 SD-Ade + 2% galactose plates. Plates were incubated for 18-20 days at 30°C. DNA libraries were sequenced (single-end, 150 bp) with custom sequencing primers on the Illumina NextSeq 500 platform. See **Supplemental Table 10** for the oligonucleotide list and Supplemental Material for more details.

## Acknowledgements

We are grateful to Grant Hartzog, Alan Hinnebusch, Fred Winston, Steven Hahn, Tim Formosa, Kevin Struhl, Craig Kaplan, Nathan Clark and John Woolford for providing yeast strains, antibodies, or plasmids. We thank Benoit Kornmann for sharing the SATAY plasmid and a data analysis script. We thank Margaret Shirra for generating integrated *spt5* substitution mutants and Tasniem Fetian for the *Os*TIR1-F74G strain used for a genetic cross. We acknowledge previous members of the lab, Mitchell Ellison, Sonia Fung, and Lydia Caldwell, for their contributions in early stages of the research. We are grateful to Sarah Hainer and Weifeng Xu for sharing equipment. We thank Fred Winston for helpful comments on the manuscript and Sarah Hainer, Craig Kaplan, Miler Lee, Carl Wu, members of their groups, and all members of the Arndt lab for helpful discussions. This project used the University of Pittsburgh Health Sciences Sequencing Core at UPMC Children’s Hospital of Pittsburgh for NGS experiments and was supported in part by the University of Pittsburgh Center for Research Computing through the resources provided. This work was supported by an Andrew Mellon Predoctoral fellowship to S.N. and NIH grant R35 GM141964 to K.M.A.

## Author Contributions

Conceptualization, S.N. and K.M.A.; funding acquisition, S.N. and K.M.A.; formal analysis, S.N.; methodology, S.N.; investigation, S.N.; validation, S.N. and K.M.A.; supervision, K.M.A.; writing—original draft, S.N. and K.M.A.; writing—review & editing, S.N. and K.M.A.

