## Supplemental Material for "Glc7/PP1 triggers Paf1 complex dissociation from RNA polymerase II to enable transcription termination"

### Contents of Supplemental Material

#### Supplemental Figures:

**Supplemental Figure S1:** Spt5 globally mediates the association of Spt6 and Paf1C with RNAPII.

**Supplemental Figure S2:** Mutations that alter the Spt5 CTR broadly affect RNAPII elongation factor occupancies and transcription at the 3’ ends of genes.

**Supplemental Figure S3:** Genes associated with RNAPII transcription regulation and RNA processing are enriched among Spt5 CTR genetic interactors.

**Supplemental Figure S4:** Mutations that disrupt the histone modification and Chd1 interaction functions of Paf1C do not rescue swd2Δ lethality.

**Supplemental Figure S5:** Dissociation of Glc7 from the 3’ ends of genes has widespread impact on RNAPII transcript synthesis.

**Supplemental Figure S6:** Glc7 anchor-away causes RNAPII elongation complex retention at the 3’ ends of genes.

**Supplemental Figure S7:** Glc7 anchor-away causes pervasive transcriptional readthrough.

#### Supplemental Materials and Methods

#### Supplemental Tables:

**Supplemental Table 1-7:** Pearson correlation values for biological replicates in ChIP-seq or 4tU-seq datasets (xlsx file).

**Supplemental Table 8:** Genotypes of yeast strains used in this study (xlsx file).

**Supplemental Table 9:** Plasmids used for yeast strain generation or experiments (xlsx file).

**Supplemental Table 10:** Oligonucleotides used for the SATAY experiments (xlsx file).

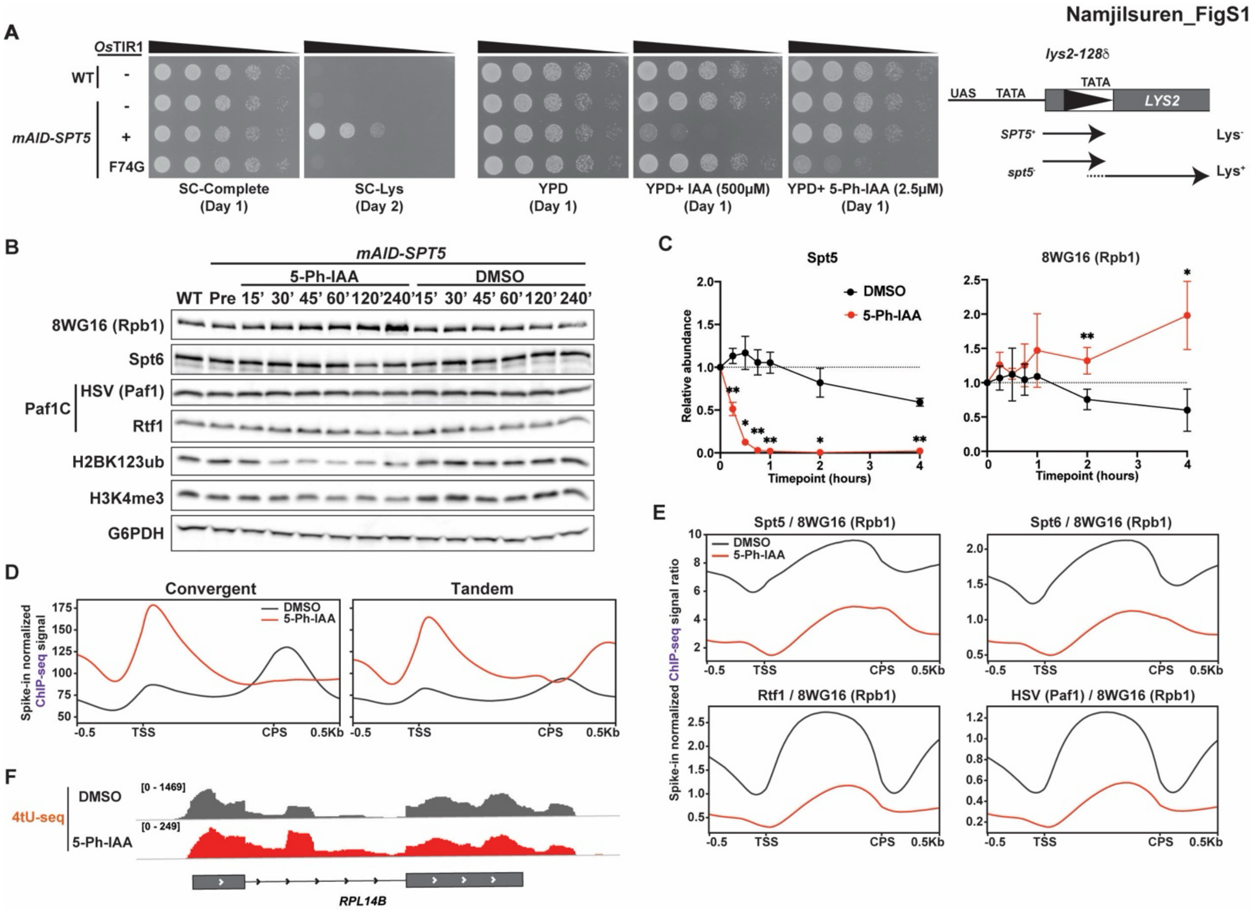


**Figure S1. Spt5 globally mediates the association of Spt6 and Paf1C with RNAPII**.

(A) Yeast spot dilution growth assay of indicated *S. cerevisiae* strains showing tight control and specificity of mAID-Spt5 degradation by the AID2 system (Yesbolatova et al. 2020). Note that with the AID2 system (*Os*TIR1-F74G) in the absence of inducer 5-Ph-IAA, the *mAID-SPT5* strain does not exhibit an Spt$^-$ phenotype with the *lys2-128∂* reporter (*i.e.* the strain exhibits a Lys$^-$ phenotype; see diagram on the right). In contrast, with the original AID system (*Os*TIR1) in the absence of inducer IAA, the *mAID-SPT5* strain is Spt$^-$ (Lys$^+$ phenotype; see diagram on the right), suggesting leaky degradation of Spt5. The diagram of *lys2-128∂* was adapted from (Swanson and Winston 1992). All subsequent experiments were performed with the AID2 system and inducer 5-Ph-IAA.

(B) Western blot analysis showing total protein abundance of elongation factors and Paf1C-dependent histone marks in the *mAID-SPT5* strain treated, over a time course, with 5-Ph-IAA or DMSO. G6PDH served as the loading control. Representative results from three independent biological replicates are shown.

(C) Quantification of western blot results from three independent replicates of experiments shown in (B) and Figure 1A. Signals are normalized to the loading control (G6PDH) and the pre-treatment condition. Mean values ± SD are shown. Significance from paired Student's *t* test with FDR correction: *$p < 0.05$ and **$p < 0.01$.

(D) Metaplots of 8WG16 (Rpb1) ChIP-seq signal for convergent (n=1981) and tandem (n=2976) genes (Hildreth et al. 2020).

(E) Metaplots of ChIP-seq signals for indicated elongation factors relative to Rpb1 signal.

(F) 4tU-seq sense strand signal at an example intron-containing gene.

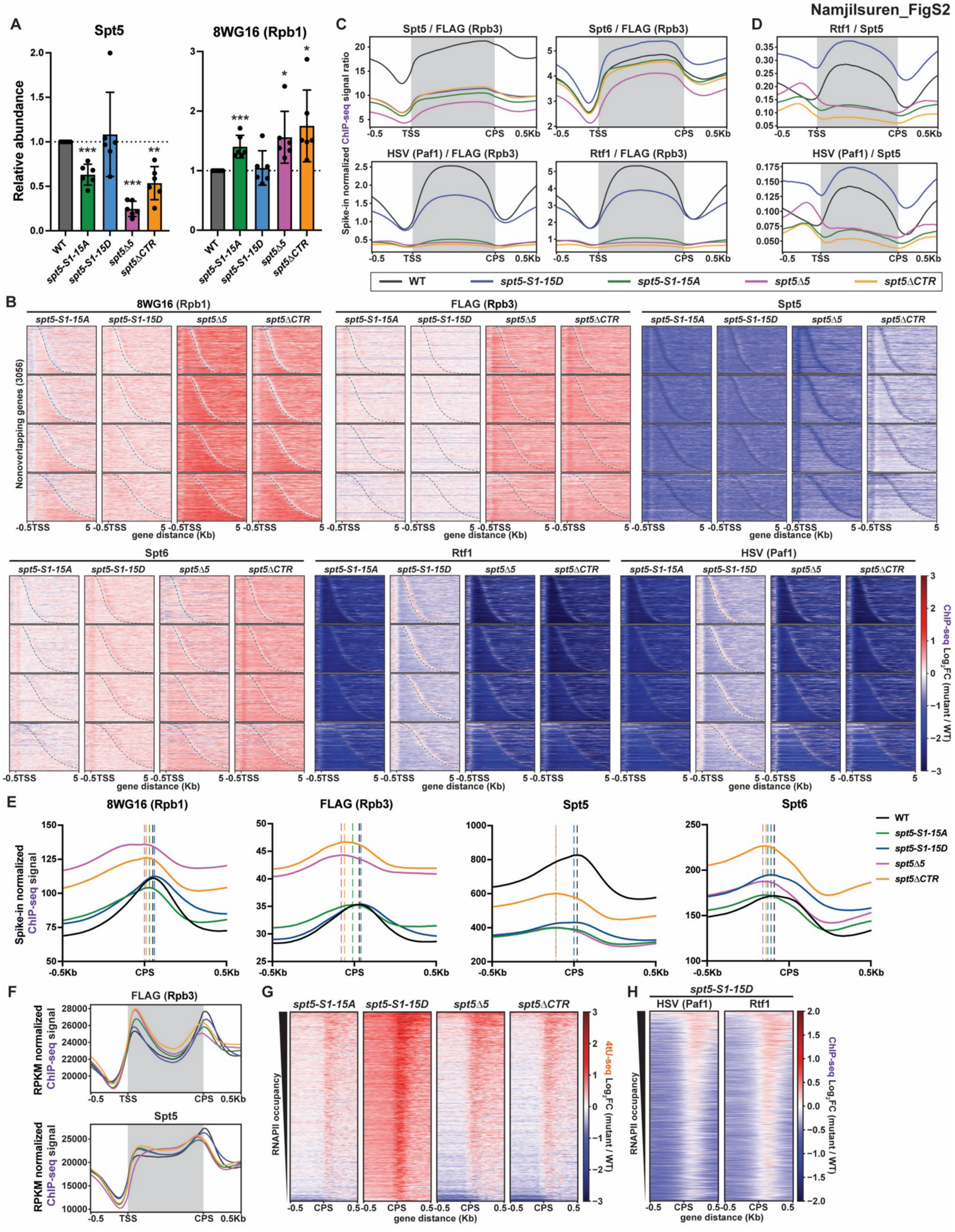


**Figure S2. Mutations that alter the Spt5 CTR broadly affect RNAPII elongation factor occupancies and transcription at the 3' ends of genes.**

(A) Quantification of western blots shown in Figure 2B. *spt5* mutants with or without Rpb3-FLAG were used for the analysis. Signals are normalized to the G6PDH loading control and to the corresponding WT. Mean values ± SD are shown. Significance from unpaired Student's *t* test: *p < 0.05, **p < 0.01 and ***p < 0.001.
(B) Differential heatmaps of ChIP-seq results for the indicated proteins and yeast strains. Genes are divided into quartiles based on Rpb1 (8WG16) occupancy for the WT strain (highest occupancy quartile on top) and sorted by gene length. Dashed line indicates CPS.
(C) Metaplots of ChIP-seq signals presented as the ratio of elongation factor occupancy relative to Rpb3-FLAG.
(D) Metaplots of Rtf1 and HSV (Paf1) ChIP-seq signals presented as a ratio relative to Spt5 ChIP-seq signal in each strain.
(E) Metaplots of ChIP-seq signals for the indicated proteins centered at the CPS. Dashed line indicates the point of maximum occupancy.
(F) Metaplots of RPKM-normalized ChIP-seq signals for the indicated proteins to highlight changes in occupancy patterns.
(G and H) Heatmaps showing $\log_2$-fold change in spike-in normalized ChIP-seq results for the indicated proteins (G) or 4tU-seq signal (H) centered at the CPS.

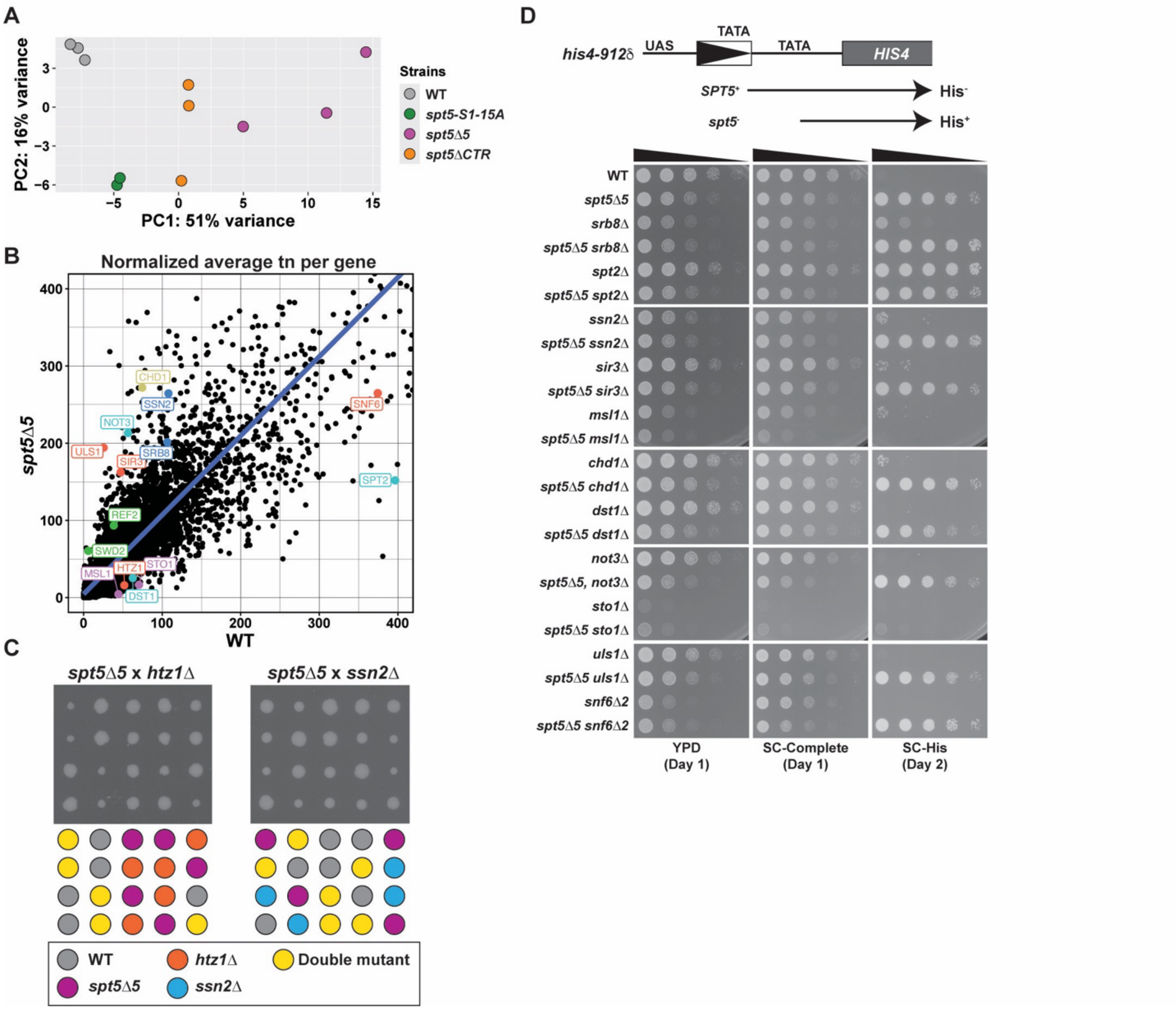


**Figure S3. Genes associated with RNAPII transcription regulation and RNA processing are enriched among Spt5 CTR genetic interactors**.

(A) PCA plot showing the correlation between transposon libraries. Technical replicates of the same biological replicate are averaged to represent a single biological replicate.

(B) Scatterplot showing normalized average transposon number per gene in the *spt5∆5* mutant and WT. Representative genes with GO terms associated with RNAPII transcription, splicing and/or chromatin organization are highlighted and were tested by *de novo* generation of deletion strains and independent genetic analysis. Genes are colored by the shared pathway.

(C) Yeast tetrad dissection plate showing examples of negative (left) or positive (right) genetic interactions with *spt5∆5*. Note that the *spt5∆5* mutation is suppressing the growth defect caused by *ssn2*∆. Each column represents four haploid spores resulting from meiosis of a single diploid. Colony growth after 3 days of growth on YPD is shown.

(D) Yeast spot dilution growth assay of single and double mutants of candidate genes shown in (B) with *spt5∆5*. Diagram explaining the Spt$^-$ phenotype for the *his4-912∂* allele is depicted (adapted from (Swanson and Winston 1992)). Growth on the SC-His plate indicates the mutant (Spt$^-$) phenotype.

Namjilsuren_FigS4

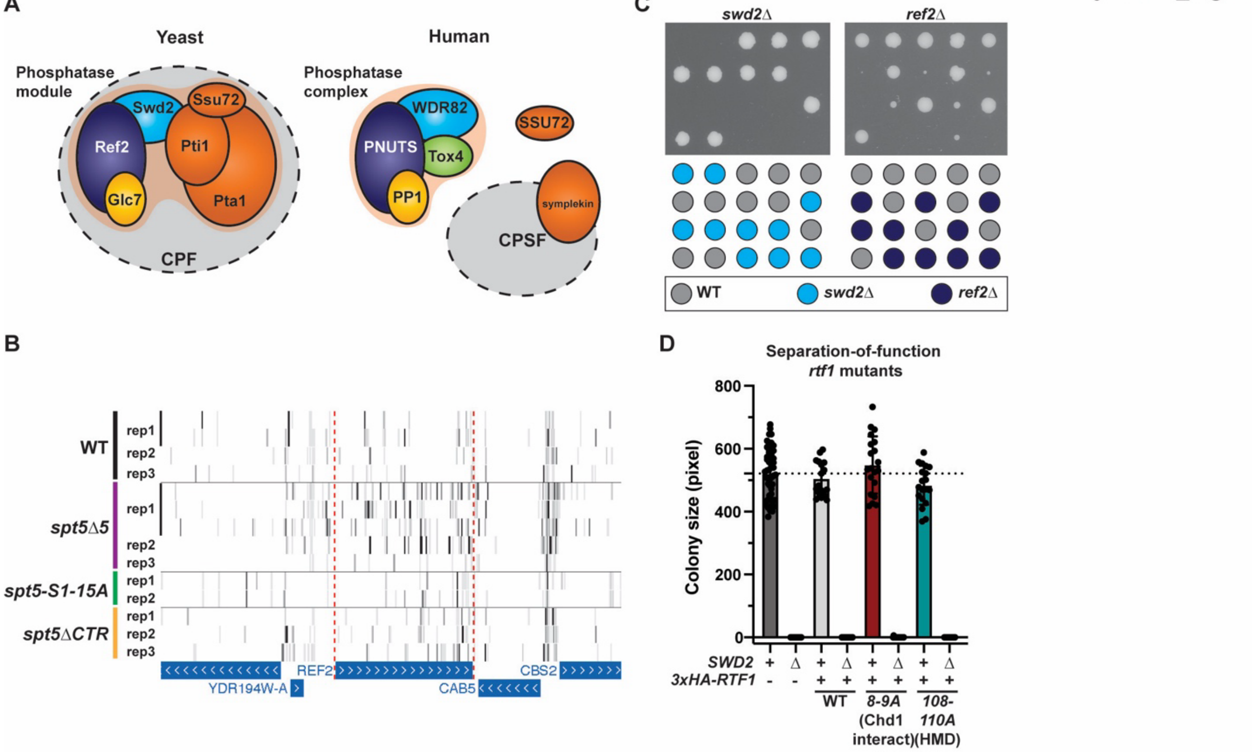


**Figure S4. Mutations that disrupt the histone modification and Chd1 interaction functions of Paf1C do not rescue *swd2*∆ lethality**.
(A) Canonical cleavage and polyadenylation (specificity) factor complexes in yeast and humans. The phosphatase module containing the Glc7 and Ssu72 phosphatases is a component of yeast CPF. In human cells, the Glc7 ortholog, PP1, is incorporated into a distinct phosphatase complex that also contains Ref2 (PNUTS) and Swd2 (WDR82) orthologs (Boreikaite and Passmore 2023).
(B) UCSC genome browser tracks showing transposon coverage in the *REF2* gene.
(C) Yeast tetrad dissection plates for heterozygous mutant diploids of *swd2*∆ and *ref2*∆.
(D) Colony size quantification of yeast tetrad dissections. Diploids heterozygous for separation-of-function *rtf1* mutations and *swd2*∆ were sporulated and progeny colonies were imaged and quantified after 3 days of growth on YPD media. Mean colony size is plotted with error bars denoting the standard deviation. Black dotted line indicates mean colony size for WT.

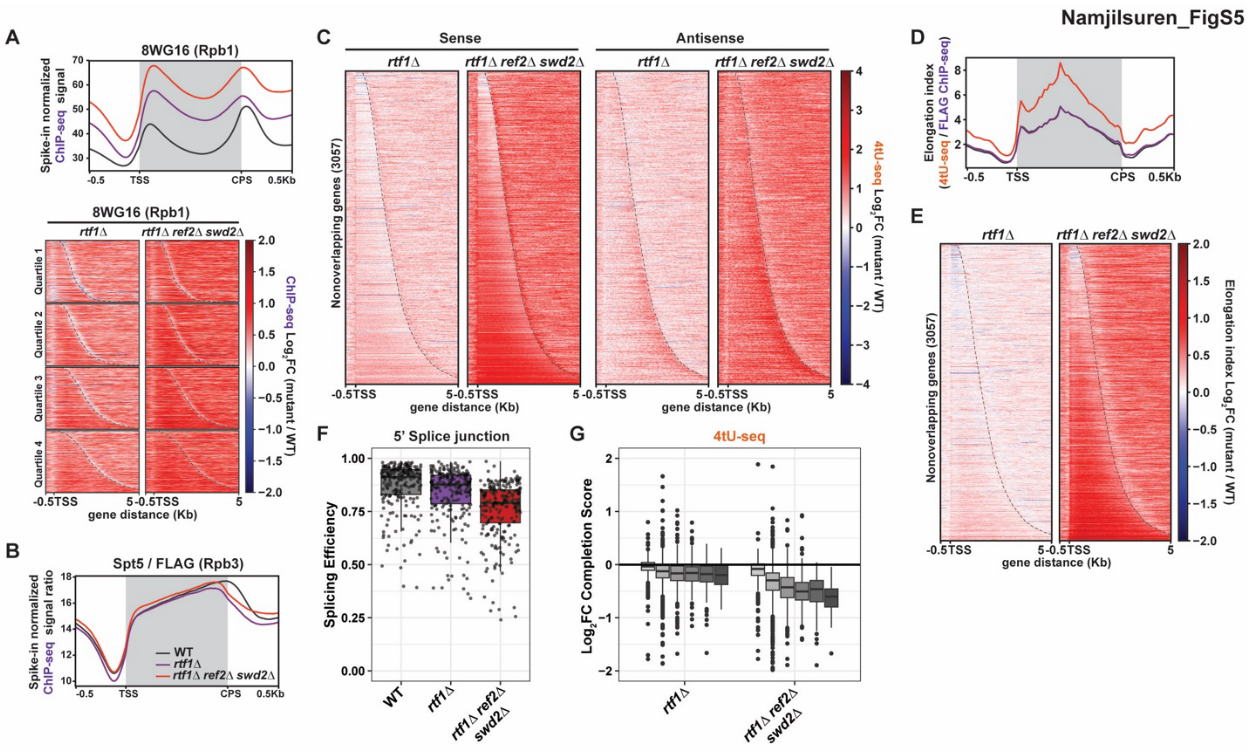


**Figure S5. Dissociation of Glc7 from the 3' ends of genes has a widespread impact on transcript synthesis**.

(A) Metaplot and heatmaps showing Rpb1 occupancy, detected with 8WG16 antibody, in the indicated mutants relative to WT. Metaplot colors are the same as in (B).

(B) Metaplot of ChIP-seq data presented as the ratio of Spt5 occupancy to RNAPII occupancy.

(C) Heatmap representation of $log_2$-fold change in 4tU-seq signal for sense and antisense transcripts.

(D and E) Metaplot (D) and differential heatmap (E) representation of elongation index.

(F-G) Boxplots report the distribution of splicing efficiency (F) and $log_2$-fold change in completion score, relative to WT, stratified by gene-length classes (G) as in Figure 2.

Namjilsuren_FigS6

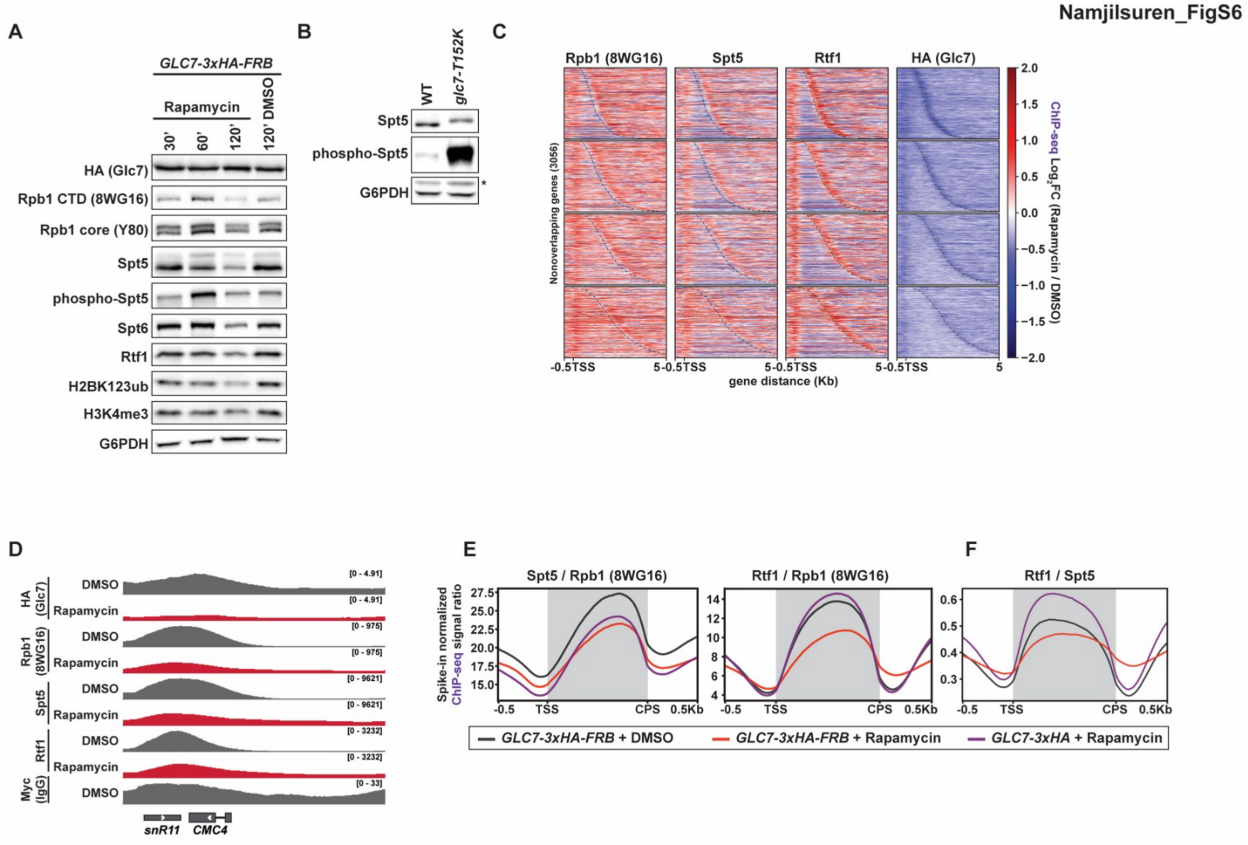


**Figure S6. Nuclear depletion of Glc7 by anchor-away causes RNAPII elongation complex retention at the 3' ends of genes**.

(A-B) Western blot analysis of Glc7 anchor-away depletion time course (A) and *glc7* mutant (B).

(C) Differential heatmaps of ChIP-seq signal for the indicated proteins in the *GLC7-3xHA-FRB* strain treated with rapamycin or DMSO for 1 hr. Genes are divided into quartiles based on Rpb1 (8WG16) occupancy (highest occupancy quartile on top) in WT (KY4803) and sorted by gene length. Dashed line indicates CPS.

(D) ChIP-seq signals at an example snoRNA locus, *snR11*.

(E-F) Metaplots of ChIP-seq data for elongation factors presented as a ratio to Rpb1 (E) or Spt5 (F) occupancy.

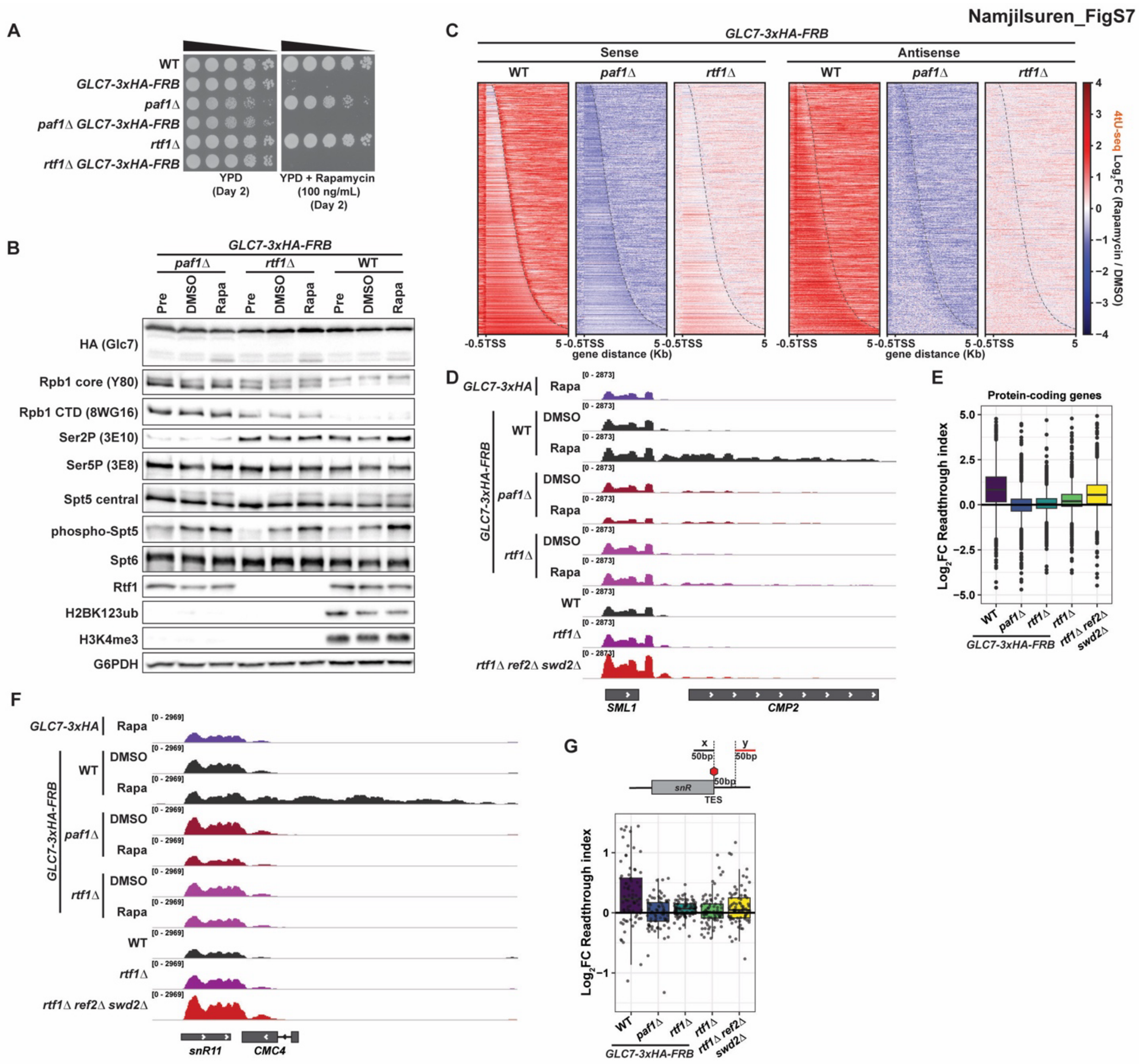


**Figure S7. Nuclear depletion of Glc7 by anchor-away causes pervasive transcriptional readthrough**.

(A) Yeast spot dilution growth assay, testing the ability of *paf1*Δ and *rtf1*Δ to suppress the growth defect on rapamycin of the Glc7 anchor-away strain.

(B) Western blot analysis of the effects of Glc7 anchor-away on the levels of various proteins and post-translational modifications in *paf1*Δ or *rtf1*Δ mutants.

(C) Heatmaps showing $\log_2$-fold change in sense and antisense transcripts upon nuclear depletion of Glc7 by anchor-away.

(D and F) Mean 4tU-seq sense-strand signal showing transcriptional readthrough at an example protein coding gene, *SML1* (D), and snoRNA locus, *snR11* (F). WT, *rtf1*Δ, and *rtf1*Δ *ref2*Δ *swd2*Δ 4tU-seq data are from Figure 5.

(E and G) Boxplots showing $\log_2$-fold change in readthrough index for non-overlapping genes (n=3057) (E) and snoRNA genes (n=77) (G). Readthrough index was calculated as in Figure 2I for protein-coding genes. For snoRNA genes, 50 bp windows, instead of 150 bp windows, were used in the calculation. Fold change reflects rapamycin/DMSO comparison for Glc7 anchor-away strains and mutant/WT comparison for the mutant strains from Figure 5.

### Supplemental Materials and Methods

*Western blot analysis and antibody information*

Protein extracts were prepared in 1x SDS-PAGE loading buffer (62.5 mM Tris-HCl pH 6.8, 2% SDS, 10% glycerol, 0.1% bromophenol blue, 5% β-mercaptoethanol (added fresh)). Proteins were resolved on 8% or 15% SDS-polyacrylamide Tris-glycine gels and transferred to nitrocellulose membranes. Nitrocellulose membranes were blocked in 5% dry milk in 1X TBST for 1 hour at room temperature. The membrane was then incubated in 5% milk in 1X TBST containing 0.02% sodium azide with primary antibody overnight at 4˚C. The primary antibodies used were: anti-Rpb1 CTD (8WG16, BioLegend, 664906, 1:500), anti-Rpb1 core (Y80, Santa Cruz, SC-25758, 1:1000), anti-Ser2P (3E10, Active Motif, 61984, 1:1000), anti-Ser5P (3E8, Active Motif, 61086, 1:500), anti-Spt5 central (gift from Grant Hartzog, 1:1000), anti-Spt6 (gift from Tim Formosa, (McCullough et al. 2015) 1:1000), anti-Rtf1 (1:2500, (Squazzo et al. 2002)), anti-HSV (Sigma-Aldrich, H6030, 1:2000), anti-FLAG (Sigma-Aldrich, F3165, 1:3000), anti-HA (Roche, 11666606001, 1:3000), anti-phospho-Spt5 (gift from Steven Hahn, 1:340, (Liu et al. 2009)), anti-H2BK123ub (Cell Signaling, 5546, 1:1000), anti-H3K4me1 (Millipore, 07-436, 1:2000), anti-H3K4me3 (Active Motif, 39159, 1:2000) and anti-G6PDH (Sigma-Aldrich, A9521, 1:20,000). The anti-Spt5 central antibody was directed against a maltose-binding protein (MBP)-Spt5 (amino acids 402-677) fusion protein. This protein was expressed from plasmid pGH79, using an MBP protein fusion and purification system from New England Biolabs (Beverly, MA), according to their instructions. Rabbits were immunized with the MBP-Spt5 fusion protein by Cocalico Biologicals (Reamstown, PA). After incubation with the primary antibody, membranes were washed with 1x TBST and incubated with secondary antibody (1:5000 dilution of HRP-conjugated anti-mouse (Cytiva, NA931) or anti-rabbit (Cytiva, NA934)) for 1 hour at room temperature. Proteins were visualized using Pico PLUS chemiluminescent substrate (Thermo Fisher, 34580) on a ChemiDoc XRS imaging platform (BioRad, Hercules, CA).

*SATAY transposon library preparation*

Briefly, 200 mL of freshly prepared SD-Ura + 2% raffinose + 0.2% dextrose media was inoculated with the *ade2*Δ *spt5* mutant yeast strains transformed with pBK549 plasmid (Michel and Kornmann 2022), which was scraped from a patch grown on a selective medium plate, at a final $OD_{600}$ of 0.16-0.18. The cultures were grown to saturation for 18 to 48 hours at 30˚C with shaking. Cells were spun down for 2 min at 1700 x g, 20˚C, and concentrated in the same media at a final $OD_{600}$ of 39. A cell suspension equivalent to an $OD_{600}$ of 7.8 was spread per plate on ~50-60 SD-Ade + 2% galactose plates using glass spreaders. Plates were incubated for 18-20 days at 30˚C. All colonies on the plates were then scraped off the plates using 200 µL of sterile water per plate and pooled. The pooled cell suspension was used to inoculate 2 L of SD-Ade + 2% dextrose culture at an initial density of $2.5 \times 10^6$ cells/mL, which was then grown to saturation at 30˚C with shaking for 24-48 hours. The saturated culture was harvested by centrifugation at 1600 x g for 5 minutes and the cell pellet was washed with water and flash frozen in liquid nitrogen as ~500 mg aliquots. Genomic DNA was extracted from a 500 mg cell pellet and 2 µg of extracted gDNA was digested with restriction enzymes *NlaIII* and *DpnII*, separately. The digested DNA was circularized using T4 ligase and then amplified by PCR with custom P5 and P7 index primers. Index 1 (i7) adapter sequences from TruSeq kits for Illumina were used as index sequences.

*ChIP-seq data processing*

ChIP-seq reads were mapped to a concatenated hybrid genome of *S. cerevisiae* (SacCer3) and *K. lactis* (ASM251v1) or *S. pombe* (ASM294v2) using bowtie2 version 2.4.5 (-p 12 -q -k 2 --no-mixed --no-discordant --no-overlap) (Langmead and Salzberg 2012). Sam files were then converted to bam files with Samtools (Li et al. 2009), and multimapping reads were filtered out with Sambamba view (-h -t 8 -F "[XS] == null" -f bam) (Tarasov et al. 2015). PCR duplicate reads were removed with Samtools markdup. Spike-in normalization factors (NormFactor) were calculated as described previously (Jeronimo et al. 2019; Ellison et al. 2023). Coverage for each base pair of the *S. cerevisiae* genome was computed using bamCoverage (--scaleFactor [NormFactor], --outFileFormat bigwig --binSize 1 --extendReads --numberOfProcessors "max") from deeptools (version 3.3.0) (Ramirez et al. 2016) to apply spike-in normalization. For RPKM normalization, --normalizeUsing RPKM option was used instead of –scaleFactor. Metaplots and heatmaps were visualized using the deeptools suite. Genes with no coverage detected in WT (KY4803) ChIP-seq datasets were removed from the non-overlapping protein-coding gene (n=3087) annotation (Reim et al. 2020) bed file and the remaining genes (n=3057) were used for all data visualization. All metaplot and heatmaps were generated from data in 10-nt bins. Individual gene signals are visualized in IGV (Robinson et al. 2011).

*4tU-seq data processing*

4tU-seq reads were first processed with Fastp (--low_complexity_filter --correction --overrepresentation_analysis) (Chen 2023) and mapped to a concatenated hybrid genome of *S. cerevisiae* (SacCer3) and *S. pombe* (ASM294v2) using HISAT2 version 2.2.1 (--rna-strandness FR --no-unal -k 2 -p 4) (Kim et al. 2019). Sam files were then converted to bam and duplicate reads were removed using the samtools suite (Li et al. 2009). *S. cerevisiae* and *S. pombe* reads were separated using Bamtools split (Barnett et al. 2011) and Samtools merge. For spike-in normalization, *S. pombe* reads over coding features were counted with featureCounts (Liao et al. 2014) and the DESeq2 estimateSizeFactors() (Love et al. 2014) function was used to generate spike-in size factors. Because the cell concentration of *S. cerevisiae* counted at the time of the harvest deviated from the number extrapolated from the linear fit by ±10%, we also applied cell count normalization to *S. cerevisiae* coverage for each base pair. The following formula was used to compute the normalization factor that takes into account both spike-in and cell count normalizations.

$$NormFactor = \frac{1}{Spike-in\ size\ factor} \times \frac{1}{\frac{(Total\ cell\ number\ of\ mutant)}{(Total\ cell\ number\ of\ WT\ (KY4803\ Rep\ 1))}}$$

Four replicates of WT Glc7 anchor-away were sequenced in total. Two replicates of WT Glc7 anchor-away were normalized post-hoc using spike-in and cell count as above. For the other two replicates of WT Glc7 anchor-away along with *paf1*Δ and *rtf1*Δ strains, only spike-in normalization was applied to *S. cerevisiae* coverage, because the total number of cells were normalized using harvested cell count information before going into RNA extraction and library build. Bam files were converted to stranded bigwig files using bamCoverage (--scaleFactor [NormFactor] --outFileFormat bigwig --binSize 1 --numberOfProcessors "max") from deeptools (version 3.3.0) (Ramirez et al. 2016). Metaplots and heatmaps were visualized in 10-nt bins using the same gene annotation set that was used for ChIP-seq and the deeptools suite. Individual gene signals are visualized in IGV (Robinson et al. 2011).

*SATAY data analysis*

The sequencing data were analyzed using a custom R script obtained from the lab of Benoit Kornmann (University of Oxford). A resulting parameter, number of transposons per gene (tnpergene), was normalized to the total number of transposons mapped in the library and multiplied by 1 million for simplicity. Normalized tn/gene data of technical replicates (separate sequencing runs for the same transformant or library) of the same biological replicate were averaged and used for all downstream analyses. Raw bed files containing all mapped transposons were uploaded to UCSC genome browser for visualization (Perez et al. 2025). Differential transposition analysis was performed using DESeq2 (Love et al. 2014) and genes that had significantly different transposon coverage were defined using the cutoff of p.adj ≤ 0.05. ClusterProfiler (Yu et al. 2012) was used to perform Gene Ontology (GO) term enrichment analysis (Gene Ontology et al. 2023).

*Elongation index*

Elongation index (Caizzi et al. 2021) was computed by taking the ratio of 4tU-seq over Pol II ChIP-seq (either Rpb3-FLAG or 8WG16) coverage in 25-nt bins using bigwigCompare from the deeptools suite (Ramirez et al. 2016). Heatmap and metaplot representations of elongation index were visualized using deeptools.

*Splicing efficiency*

4tU-seq total read coverage (including both spliced and unspliced) overlapping +1 base (5' splice site) (Prevorovsky et al. 2016; Francette and Arndt 2024) of each annotated chromosomal intron feature from *S. cerevisiae* gff file from SGD (Wong et al. 2023) was determined using bedtools coverage (-s -a -counts) (Quinlan and Hall 2010). Unspliced read coverage was determined with an additional -split option. Introns with spliced read coverage below an arbitrary threshold of 5 were excluded to avoid artifacts originating from low gene expression level. Splicing efficiency was calculated by dividing the read coverage of spliced reads by the total read coverage of spliced and unspliced reads per protein-coding gene intron.

*Completion score*

4tU-seq read density overlapping 200 bp windows, positioned 100 bp downstream of the TSS (proximal) or upstream of the CPS (distal), for the non-overlapping gene set (described above) was determined using multiBigwigSummary BED-file command. Only reads aligning on the sense strand were considered. Completion score was calculated by dividing distal read coverage by proximal read coverage and computing the $\log_2$ fold change (Narain et al. 2021; Francette and Arndt 2024).

*Readthrough index*

4tU-seq read density overlapping 150 bp (50 bp for snoRNAs) windows immediately upstream (pre-CPS) or downstream (readthrough) of the CPS of the non-overlapping gene set was computed by multiBigwigSummary BED-file command. Readthrough index was calculated as the $\log_2$ fold change of readthrough read density compared to pre-CPS signal (Narain et al. 2021).

*Statistical analysis and data reproducibility*

All genomic experiments were performed in biological duplicate, except for 4tU-seq analysis of the WT Glc7 anchor-away strain for which four replicates were included in Fig. 7. Pearson's correlation analysis was used to assess replicate reproducibility. Each biological replicate is a yeast culture derived from a single colony. For calculations of splicing efficiency, completion score and readthrough index, *p*-values were generated using Mann-Whitney U test (wilcox.test in R) with FDR correction. For SATAY, at least two biological replicates (independent transformants) with 1-3 technical replicates were analyzed for all strains. Principal component analysis was used to assess SATAY replicate reproducibility. Quantification data were visualized using ggplot2 (Wickham 2011). For yeast tetrad dissections, at least two independently generated diploid strains were assessed. Western blots were performed in biological triplicate or more, except for *glc7-T152K* mutant which had two replicates, and *p*-values associated with the quantification were calculated in GraphPad Prism as indicated in the figure legends.
